# Novel lentiviral vectors for gene therapy of sickle cell disease combining gene addition and gene silencing strategies

**DOI:** 10.1101/2022.12.31.522279

**Authors:** Mégane Brusson, Anne Chalumeau, Pierre Martinucci, Oriana Romano, Valentina Poletti, Samantha Scaramuzza, Sophie Ramadier, Cecile Masson, Giuliana Ferrari, Fulvio Mavilio, Marina Cavazzana, Mario Amendola, Annarita Miccio

**Author notes:** To whom correspondence should be addressed: M.B., Imagine Institute, 24, Boulevard du Montparnasse, 75015 Paris, France. E-mail address. A.M., Imagine Institute, 24, Boulevard du Montparnasse, 75015 Paris, France.

## Abstract

Sickle cell disease (SCD) is due to a mutation in the β-globin (*HBB*) gene causing the production of the toxic sickle hemoglobin (HbS, a_2_β^S^_2_). Transplantation of autologous hematopoietic stem/progenitor cells (HSPCs) transduced with lentiviral vectors (LVs) expressing an anti-sickling β-globin (βAS) is a promising treatment; however, it is only partially effective and patients still present elevated HbS levels. Here, we developed a bifunctional LV expressing βAS3-globin and an artificial microRNA (amiR) specifically downregulating β^S^-globin expression with the aim of reducing HbS levels and favoring βAS3 incorporation into Hb tetramers. Efficient transduction of SCD HSPC by the bifunctional LV led to a substantial decrease of β^S^-globin transcripts in HSPC-derived erythroid cells, a significant reduction of HbS^+^ red cells and effective correction of the sickling phenotype, outperforming βAS gene addition and *BCL11A* gene silencing strategies. The bifunctional LV showed a standard integration profile and neither the HSPC viability, engraftment and multi-lineage differentiation nor the erythroid transcriptome and miRNAome were affected by the treatment, confirming the safety of this therapeutic strategy. In conclusion, the combination of gene addition and gene silencing strategies can improve the efficacy of current LV-based therapeutic approaches without increasing the mutagenic vector load, thus representing a novel treatment for SCD.

## Introduction

Mutations in the adult β-globin (*HBB*) locus lead to abnormal production of the β-globin chain of the adult Hb (HbA, α_2_β_2_) tetramers, causing β-hemoglobinopathies, the most common monogenic disorders worldwide. In β-thalassemia, reduced (β^+^) or absent (β^0^) β-globin chain production is responsible for precipitation of uncoupled α-globin chains, which in turn leads to apoptosis of erythroid precursors and impairment in erythroid differentiation (i.e. ineffective erythropoiesis), and hemolytic anemia (Thein, 2018). Sickle cell disease (SCD) is due to the production of a mutant β^S^-globin forming the sickle Hb (HbS, α_2_β^S^_2_), which has the propensity to polymerize under deoxygenated conditions, resulting in the production of sickle-shaped red blood cells (RBCs). Sickle RBCs cause occlusions of small blood vessels, leading to impaired oxygen delivery to tissues, multi-organ damage, severe pain and early mortality (Piel et al., 2017).

Transplantation of autologous hematopoietic stem and progenitor cells (HSPCs) corrected by lentiviral (LV) vectors expressing a transgene-encoded β-globin is a promising therapeutic option. However, gene addition strategies are at best partially effective in correcting the clinical phenotype in patients with severe β-thalassemia (e.g., β^0^/β^0^ patients with no residual expression of the β-globin gene) and SCD (where high expression of the β^S^-globin impairs the incorporation of the therapeutic β-globin transgene into Hb) (Cavazzana et al., 2019; Kanter et al., 2021; Locatelli et al., 2021; Magrin et al., 2022, 2019; Ribeil et al., 2017). Total Hb levels were < 12 g/dl for the vast majority of patients, indicating death of uncorrected RBCs, and HbS levels remain high (representing >50% of the total Hb) (Kanter et al., 2021; Magrin et al., 2022). On the contrary, asymptomatic SCD carriers have only 30-40% of HbS and HbS levels are usually maintained below 30% (though RBC transfusion) in severe SCD patients to suppress SCD symptoms (Ware et al., 2017). However, the distribution of therapeutic Hb and HbS within the RBCs is also an important determinant of the clinical outcome of SCD patients. By way of example, with HbF representing 20% of the total Hb, the proportion of total RBCs that expressed sufficient therapeutic Hb levels (~10 pg of HbF per RBC) to inhibit HbS polymerization varies between 1% and 24% (Steinberg et al., 2014). Almost 30% of HbF is required to have 70% of HbF^+^ RBCs protected from Hb polymerization (Steinberg et al., 2014).

Moreover, in gene addition strategies, the high number of integration sites per cell required to express clinically relevant levels of the β-globin transgene is often limited by the poor LV transduction efficiency in HSPCs or can increase the potential genotoxic risks associated with integrating LVs (Cavazzana et al., 2019; Cavazzana-Calvo et al., 2010; Goyal et al., 2021).

We have developed a high-titer LV carrying a potent anti-sickling β-globin transgene (βAS3; anti-sickling amino-acid substitutions: G16D for conferring to the transgene a competitive advantage over the sickle β-globin in terms of the interaction with the a-globin polypeptide, E22A for disrupting axial contacts in Hb polymers, T87Q for blocking the lateral contact with valine 6 of the sickle β-globin chain) under the control of *HBB* promoter and a mini-locus control region (LCR) (Weber et al., 2018). However, despite the good gene transfer efficiency in SCD HSPCs, RBC sickling was only partially corrected, even at a high vector copy number per cell (VCN/cell), due to the residual HbS expression.

The clinical severity of SCD and β-thalassemia is alleviated by the co-inheritance of genetic mutations causing a sustained fetal γ-globin (*HBG1* and *HBG2*) chain production at adult age, a condition termed hereditary persistence of fetal Hb (HPFH). SCD-HPFH patients with fetal Hb (HbF, α_2_γ_2_) representing >20% of the total Hb show a less severe disease phenotype and improved survival (Powars et al., 1984). Gamma-globin exerts an anti-sickling effect in SCD by displacing the β^S^-globin from the Hb tetramer to form HbF or mixed Hb tetramers (α_2_βγ) that cannot polymerize (Akinsheye et al., 2011). Gamma-globin reactivation has been extensively explored as a therapeutic approach for β-hemoglobinopathies. A gene silencing strategy based on a LV expressing an artificial microRNA (amiR), consisting of a shRNA embedded in a miR backbone (shmiR), has been exploited to specifically downregulate the *BCL11A* extra-large (XL) transcript (*BCL11A-XL*) encoding the BCL11A isoform responsible for γ-globin repression in adulthood (Brendel et al., 2016; Guda et al., 2015). The therapeutic benefit of HbF reactivation using LV strategies is currently under evaluation in a clinical trial (Brendel et al., 2020; Esrick et al., 2021). Despite the early promising clinical data, HbF levels were only ~25% to ~40%, HbS levels were modestly reduced, and Hb concentration was still < 12 g/dl (Esrick et al., 2021).

Therefore, despite the undeniable progress in the field of gene therapy for β-hemoglobinopathies, additional improvements in LV design and LV-based gene therapy are required to obtain a robust clinical benefit in severe β-thalassemia and SCD.

In this study, we generated two novel LV vectors combining a βAS3 gene addition approach with an amiR-based gene silencing strategy aimed at down-regulating either BCL11A (βAS3/miRBCL11A) or the β^S^-globin (βAS3/miRHBB), to further boost therapeutic β-like globin levels without increasing the mutagenic vector load in HSPCs and correct the β-hemoglobinopathy phenotype. By downregulating BCL11A, amiRBCL11A re-activated the expression of the endogenous anti-sickling fetal γ-globin in erythroid cells differentiated from β-thalassemia and SCD HSPCs. However, in SCD RBCs, HbF induction was not associated with a concomitant reduction of HbS expression levels required to correct the disease phenotype. On the contrary, the therapeutic strategy based on β^S^-globin downregulation combined with βAS3 expression led to a strong reduction of HbS levels and HbS^+^ RBCs derived from SCD HSPCs, which favored βAS3 incorporation in Hb tetramers, increased therapeutic Hb levels and ameliorated the SCD phenotype

## Materials and Methods

### Generation of lentiviral constructs

The pCCL.b-AS3 plasmid (βAS3 LV) was used to generate all LV constructs used in this study (Weber et al., 2018). The *HBB* mini-gene (coding for the βAS3 transgene) contains three mutations determining three amino acid substitutions (G16D, E22A, and T87Q) and a 593-bp deletion in intron 2 removing a region located 85 to 679 bp downstream of *HBB* exon 2 (total length of intron 2 is 257 bp). We inserted the miRBCL11A between positions 85 and 86 (hereafter named Int2_del) or between 146 and 147 (hereafter named Int2) of the βAS3 intron 2. We did not test additional positions as we have a short intron and we needed to avoid positions that are too close to the splicing junctions. We inserted the different shRNA-, siRNA- or miR-derived sequences in the miR223 backbone (Amendola et al., 2009) and cloned the miR in βAS3 intron 2. The guide strand of the miRBCL11A was previously published (Brendel et al., 2016), while those targeting *HBB* are displayed in **Table 1**. In particular, for miRs derived from shRNA, the sequence was either used as such or modified by adding a GCGC motif at the 3’ end of the guide strand and removing the 4 nucleotides at the 5’ end (**Table 1**) (Guda et al., 2015). We used a non-targeting miR (miRnt) as control; it targets the LacZ mRNA, (5′-AAATCGCTGATTTGTGTAGTC-3′; Amendola et al., 2009). In the LV βAS3m/miR7m, containing the most efficient miRHBB, the βAS3 transgene was modified by inserting synonymous mutations as described in **Figure 5**.

**Table 1:**
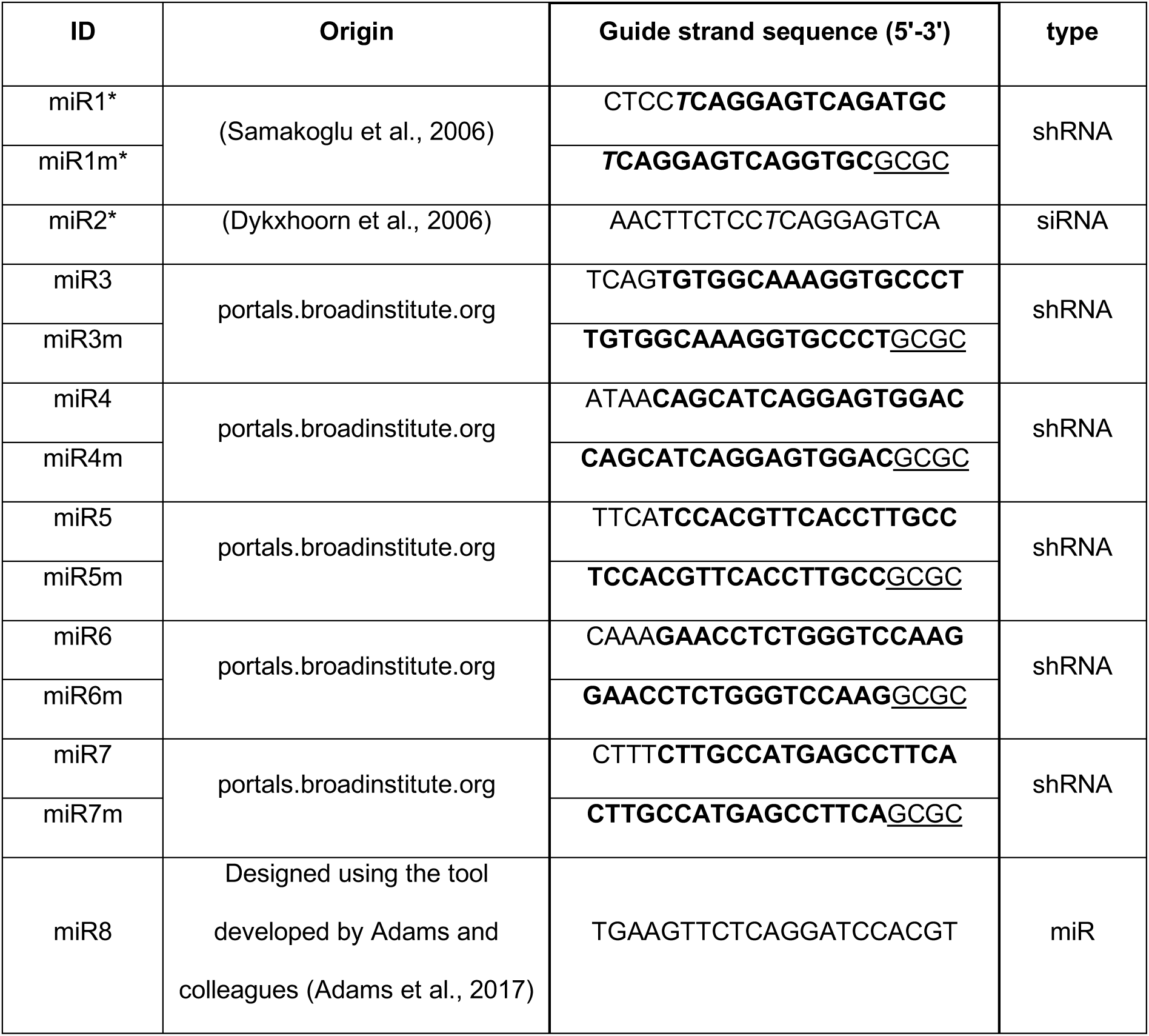
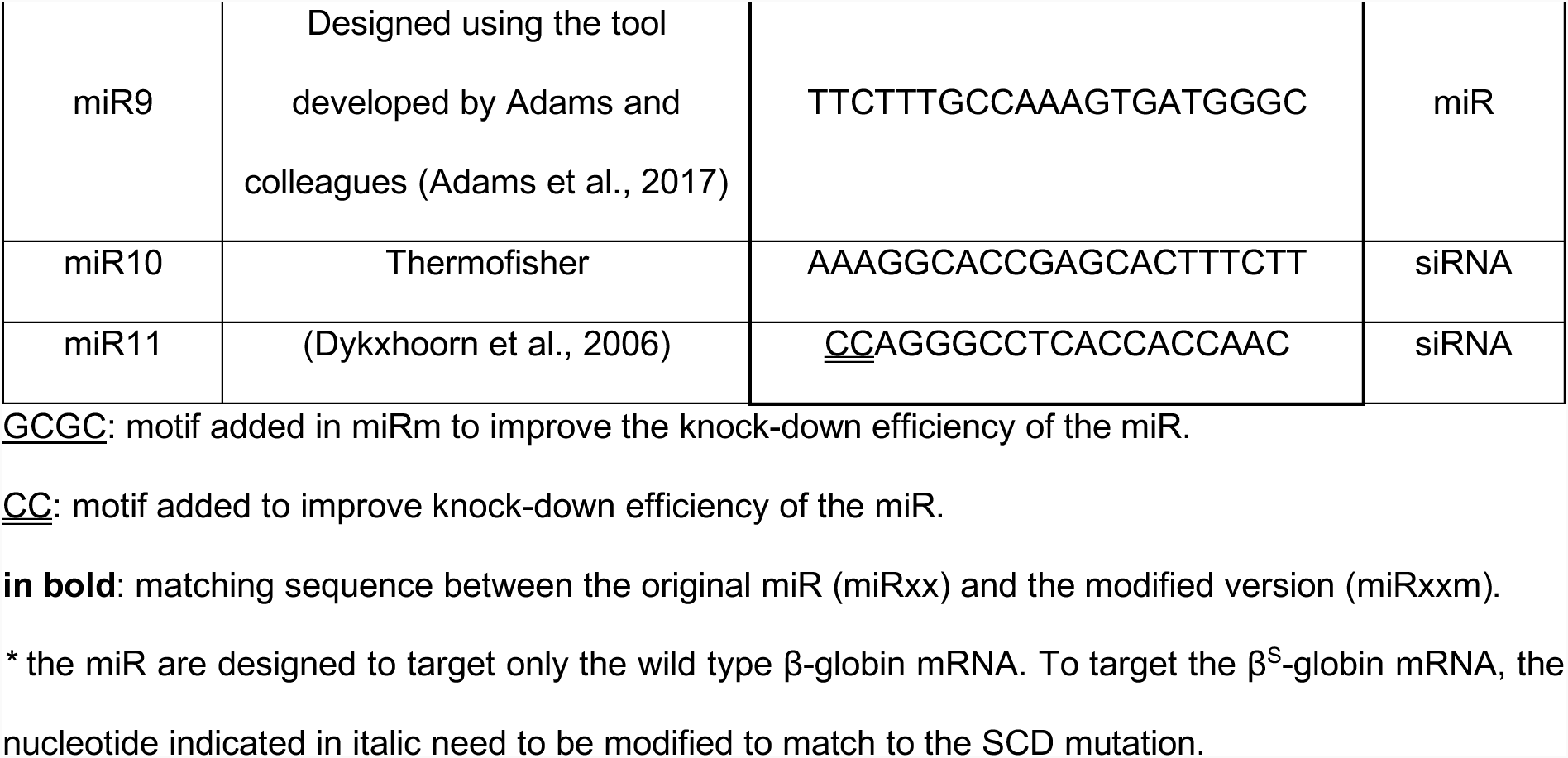
Guide strand sequences targeting the *HBB* mRNA.

### Lentiviral vector production and titration

Third-generation LVs were produced by calcium phosphate-based transient transfection of HEK293T cells with the transfer vector, the packaging plasmid pHDMH gpm2 (encoding gag/pol), the Rev-encoding plasmid pBA Rev, and the vesicular stomatitis virus glycoprotein G (VSV-G) envelope-encoding plasmid pHDM-G. The physical titer of vector preparations was measured using the HIV-1 Gag p24 antigen immunocapture assay kit (PerkinElmer, Waltham, MA, USA) and expressed as p24 ng/ml. The viral infectious titer, expressed as transduction units per ml (TU/ml) was measured in HCT116 cells after transduction using serial vector dilutions. VCN/cell was measured by ddPCR. The LV titer was then calculated as follows: Titer (TU/ml) = (number of transduced cells*VCN)/volume of vector used. Viral infectivity was calculated as ratio between infectious and physical titers (TU/ng p24).

### HUDEP-2 cell culture, differentiation and transduction

HUDEP-2 cells were cultured and differentiated as previously described (Antoniani et al., 2018; Canver et al., 2015; Kurita et al., 2013). HUDEP-2 cells were expanded in a basal medium composed of StemSpan SFEM (Stem Cell Technologies) supplemented with 10^−6^ M dexamethasone (Sigma), 100 ng/ml human stem cell factor (hSCF) (Peprotech), 3 IU/ml erythropoietin (EPO) Eprex (Janssen-Cilag, France), 100 U/ml L-glutamine (Life Technologies), 2 mM penicillin/streptomycin and 1 μg/ml doxycycline (Sigma). HUDEP-2 cells were transduced at a cell concentration of 10^6^ cells/ml in basal medium supplemented with 4 μg/ml protamine sulfate (Choay). After 24 h, cells were washed and cultured in fresh basal medium. Cells were differentiated for 9 days in Iscove’s Modified Dulbecco’s Medium (IMDM; Life Technologies) supplemented with 330 μg/ml holo-transferrin (Sigma), 10 μg/ml recombinant human insulin (Sigma), 2 IU/ml heparin (Sigma), 5% human AB serum (Eurobio AbCys), 3 IU/mL erythropoietin, 100 ng/mL hSCF, 1 μg/ml doxycycline, 100 U/ml L-glutamine, and 2 mM penicillin/streptomycin.

### K562 cell culture and transduction

K562 cells were maintained in RPMI 1640 medium (Lonza) containing glutamine and supplemented with 10% fetal bovine serum (Lonza), HEPES (LifeTechnologies), sodium pyruvate (LifeTechnologies) and penicillin/streptomycin (LifeTechnologies). K562 cells were transduced at a cell concentration of 5×10^5^ cells/ml in the culture medium supplemented with 4 μg/ml polybrene (Sigma). After 24 h, cells were washed and cultured in fresh medium.

### HSPC purification, culture and transduction

Peripheral blood plerixafor-mobilized or non-mobilized human adult HSPCs were obtained from SCD patients. Peripheral blood G-CSF (granulocyte colony stimulating factor)-mobilized human adult HSPCs were obtained from healthy donors (HD). Written informed consent was obtained from all subjects. All experiments were performed in accordance with the Declaration of Helsinki. The study was approved by the regional investigational review board (reference DC 2014-2272, CPP Ile-de-France II “Hôpital Necker-Enfants malades”; Paris, France). SCD and HD HSPCs were purified by immunomagnetic selection (Miltenyi Biotec) after immunostaining using the CD34 MicroBead Kit (Miltenyi Biotec). Plerixafor/G-CSF-mobilized peripheral blood CD34^+^ cells were selected from patients affected by β-thalassemia upon signed informed consent approved by the Ethical Committee of the San Raffaele Hospital (Milan, Italy). Following mobilization and cell collection with the Spectra Cobe or Spectra Optia apheresis system (Terumo BCT), CD34^+^ cells were purified using immunomagnetic beads (CliniMACS, Miltenyi Biotec) by MolMed SpA (Milan, Italy) (Marktel et al., 2019).

SCD and HD CD34^+^ cells were thawed and cultured for 24 h at a concentration of 10^6^ cells/mL in a pre-activation medium (PAM) composed of X-VIVO 20 supplemented with penicillin/streptomycin (Gibco) and recombinant human cytokines: 300 ng/mL hSCF, 300 ng/mL Flt-3L, 100 ng/mL TPO, 20 ng/mL interleukin-3 (IL-3) (Peprotech) and 10 mM SR1 (StemCell). After pre-activation, cells (10^6^ cells/mL) were cultured in PAM supplemented with 10 μM PGE2 (Cayman Chemical) on RetroNectin-coated plates (10 μg/cm^2^, Takara Bio) for at least 2 h. Cells (10^6^ cells/mL) were then transduced for 24 h on RetroNectin-coated plates in PAM supplemented with 10 μM PGE2, protamine sulfate (4 μg/mL, Protamine Choay) and Lentiboost (1 mg/ml, SirionBiotech). β-thalassemia CD34^+^ cells were thawed and cultured for 24 h at a concentration of 10^6^ cells/mL in pre-activation medium composed of CellGro (Corning) supplemented with 300 ng/mL hSCF, 300 ng/mL Flt-3L, 100 ng/mL TPO, 20 ng/mL IL-3 (Peprotech) on RetroNectin-coated plates (10 μg/cm^2^, Takara Bio). Cells were then transduced in the same medium, supplemented with 10 μM PGE2 (Cayman Chemical) and Lentiboost (1 mg/ml, SirionBiotech) for 18 hours in presence of LV vector at the indicated MOI.

### Vector copy number quantification by ddPCR

Genomic DNA was extracted from HCT116 cells 4 days after transduction, and for K562 cells, HUDEP-2 cells, primary erythroblasts, burst-forming unit erythroid (BFU-E), colony-forming unit for granulocytes and macrophages (CFU-GM) colonies 14 days after transduction, and human CD45^+^ (hCD45^+^) bone marrow (BM) cells from transplanted mice using the PureLink Genomic DNA Mini Kit (Invitrogen). DNA was digested using DraI restriction enzyme (NEB) at 37°C for 30 min and then mixed with the ddPCR reaction mix composed of 2X ddPCR SuperMix for probes (no dUTP) (Bio-Rad), forward (for) and reverse (rev) primers (at a final concentration of 900 nM) and probes (at a final concentration of 250 nM). We used probes and primers specific for: (i) albumin (VIC-labeled ALB probe with a QSY quencher, 5’-CCTGTCATGCCCACACAAATCTCTCC-3’; FOR ALB primer, 5’-GCTGTCATCTCTTGTGGGCTGT-3’; REV ALB primer, 5’-ACTCATGGGAGCTGCTGGTTC-3’), and for (ii) the LV (FAM-labeled LV probe with an MGB quencher, 5’-CGCACGGCAAGAGGCGAGG-3’; FOR LV primer 5’-TCCCCCGCTTAATACTGACG-3’; REV LV primer 5’-CAGGACTCGGCTTGCTGAAG-3’ or FAM-labeled PRO-LV probe with an MGB quencher 5’-TCTCTAGCAGTGGCGCCCGAACAGG-3’; FOR PRO-LV primer: 5’-CACTCCCAACGAAGACAAGA-3’; REV PRO-LV primer: 5’-TCTGGTTTCCCTTTCGCTTT-3’ to measure VCN in BM cells (Corre et al., 2022). The albumin gene was chosen as reference locus to calculate the VCN per genome. Droplets were generated using a QX200 droplet generator (Bio-Rad) with droplet generation oil for probes (Bio-Rad) onto a DG8 cartridge (Bio-Rad) and transferred on a semi-skirted 96 well plate (Eppendorf AG). After sealing with a pierceable foil heat seal using a PX1 PCR plate sealer (Bio-Rad), the plate was loaded on a SimpliAmp Thermal Cycler (ThermoFisher Scientific) for PCR amplification using the following conditions: 95°C for 10 min, followed by 40 cycles at 94°C for 30 sec and 60°C for 1 min, and by a final step at 98°C for 10 min. The plate was analyzed using the QX200 droplet reader (Bio-Rad) (channel 1: FAM, channel 2: VIC) and analyzed using the QuantaSoft analysis software (Bio-Rad), which quantifies positive and negative droplets and calculates the starting DNA concentration using a Poisson algorithm. The average VCN per cell was calculated as (LV copies*x)/(albumin copies), where x=2 for diploid cells and 4 for tetraploid cells.

To evaluate transduction efficiency in colony-forming cell (CFC)-derived progenitors from β-thalassemia patients, single BFU-E and CFU-GM were lysed with QuickExtract Lysis Buffer (Epicentre). CFCs were incubated at 65°C for 20 min followed by an incubation at 98°C for 10 min and centrifuged at 13000 rpm for 10 minutes. VCN was evaluated by ddPCR using the previously described HIV/Telo system (Marktel et al., 2019).

### *In vitro* erythroid differentiation

Mature RBCs from mock- and LV-transduced CD34^+^ HSPCs were generated using a three-step protocol (Weber et al., 2018). Briefly, from day 0 to 6, cells were grown in a basal erythroid medium (BEM) supplemented with hSCF, IL-3, EPO (Eprex, Janssen-Cilag) and hydrocortisone (Sigma). From day 6 to 20, cells were cultured on a layer of murine stromal MS-5 cells in BEM supplemented with EPO from day 6 to day 9, and without cytokines from day 9 to day 20. From day 13 to 20, human AB serum was added to the BEM.

### CFC assay

The number of hematopoietic progenitors was evaluated using a colony-forming cell (CFC) assay. HSPCs were plated at a concentration of 5×10^2^ cells/mL in a methylcellulose-containing medium (GFH4435, Stem Cell Technologies) under conditions supporting erythroid and granulo-monocytic differentiation. BFU-E and CFU-GM colonies were scored after 14 days. BFU-E and CFU-GM were randomly picked and collected as bulk populations (containing at least 25 colonies) to evaluate transduction efficiency and globin expression.

### RT-qPCR analysis

RNA was extracted from HUDEP-2 cells, K562 cells, primary erythroblasts or BFU-E using the RNeasy micro kit (QIAGEN). Reverse transcription of mRNA was performed using the SuperScriptIII First-Strand Synthesis System for RT-PCR (Invitrogen) with oligo(dT)_20_ primers. qPCR was performed using the SYBR green detection system (BioRad). We used the following primers: HBB FOR, 5’-AAGGGCACCTTTGCCACA-3’; HBB REV, 5’-GCCACCACTTTCTGATAGGCAG-3’; βAS3 FOR, 5’-GCCACCACTTTCTGATAGGCAG-3’; βAS3 REV, 5’-AAGGGCACCTTTGCCCAG-3’; BCL11A-XL FOR, 5’-ATGCGAGCTGTGCAACTATG-3’; BCL11A-XL REV, 5’-GTAAACGTCCTTCCCCACCT-3’; HBG1/2 FOR, 5’-CCTGTCCTCTGCCTCTGCC-3’; HBG1/2 REV, 5’-GGATTGCCAAAACGGTCAC-3’; LMNB2 FOR, 5’-AGTTCACGCCCAAGTACATC-3’; LMNB2 REV, 5’-CTTCACAGTCCTCATGGCC-3’; HBA FOR, 5’-CGGTCAACTTCAAGCTCCTAA-3’; HBA REV, 5’-ACAGAAGCCAGGAACTTGTC-3’; GAPDH FOR, 5’-GAAGGTGAAGGTCGGAGT-3’; GAPDH REV, 5’-GAAGATGGTGATGGGATTTC-3’. The samples were analyzed with the ViiA 7 Real-Time PCR System and software (Applied Biosystems).

### Flow cytometry analysis

After 9 days of differentiation, HUDEP-2 cells were stained with a monoclonal mouse anti-human CD235a antibody (clone GA-R2, BD Biosciences), then fixed and permeabilized with the fixation/permeabilization solution kit (BD Biosciences), and stained with a monoclonal mouse anti-human HbF antibody (clone HBF-1, ThermoFisher scientific). Cells were analyzed by flow cytometry using a BD LSRFortessa cell analyzer (BD Biosciences) and the Diva (BD Biosciences) and the FlowJo software.

In primary cell cultures, the expression of erythroid markers was monitored by flow cytometry using anti-CD36, anti-CD49d, anti-CD71, anti-CD235a (BD Biosciences) and anti-CD233 (band 3; IBGRL) antibodies and 7-AAD (BD Biosciences) for cell death assessment. The proportion of enucleated RBCs was measured using the nuclear dye DRAQ5 (eBioscience). The proportion of HbF-and HbS-positive RBCs were measured with an antibody recognizing HbF (Thermo Fisher for HUDEP-2 cells or BD Biosciences for primary cells) or HbS (BioMedomics), respectively. Briefly, RBCs were stained with a monoclonal mouse anti-human CD235a antibody (BD Biosciences), then fixed with 0.05% glutaraldehyde for 10 min at RT, permeabilized with 0.1% Triton X-100 for 10 min at RT, and stained with the HbF or the HbS antibody. Flow cytometry analyses were performed using the Gallios analyzer and Kaluza (Beckman-Coulter) and the FlowJo software.

### Western blot

HUDEP-2 cells after 6 days of differentiation, to detect BCL11A-XL, were lysed for 30 min at 4°C using a lysis buffer containing: 10 mM Tris, 1 mM EDTA, 0.5 mM EGTA, 1% Triton X-100, 0.1% SDS, 0.1% Na-deoxicholate, 140 mM NaCl (Sigma-Aldrich) and a protease inhibitor cocktail (Roche-Diagnostics). Cell lysates were sonicated twice (50% amplitude, 10 sec per cycle, pulse 9 sec on/1 sec off) and underwent 3 cycles of freezing/thawing (3 min at −80°C/3 min at 37°C). After centrifugation, the supernatant was collected and protein concentration was measured using the Pierce™ BCA Protein Assay Kit (ThermoScientific). After electrophoresis and protein transfer, BCL11A-XL, and GAPDH were detected using the antibodies ab19487 (abcam) and sc-32233 (SantaCruz), respectively. The bands corresponding to the different proteins were quantified using the Chemidoc and the Image lab Software (BioRad).

### HPLC

High-performance liquid chromatography (HPLC) analysis was performed using a NexeraX2 SIL-30AC chromatograph (Shimadzu) and the LC Solution software. Globin chains from differentiated HUDEP-2 cells, primary cells, or BFU-E were separated by reverse-phase (RP)-HPLC using a 250×4.6 mm, 3.6 µm Aeris Widepore column (Phenomenex). Samples were eluted with a gradient mixture of solution A (water/acetonitrile/trifluoroacetic acid, 95:5:0.1) and solution B (water/acetonitrile/trifluoroacetic acid, 5:95:0.1). The absorbance was measured at 220 nm. Hb tetramers from RBCs were separated by cation exchange (CE)-HPLC using a 2 cation-exchange column (PolyCAT A, PolyLC, Columbia). Samples were eluted with a gradient mixture of solution A (20mM bis Tris, 2mM KCN, pH, 6.5) and solution B (20mM bis Tris, 2mM KCN, 250mM NaCl, pH, 6.8). The absorbance was measured at 415nm.

### Sickling assay

At day 19 of the terminal erythroid differentiation, RBCs were collected and incubated under hypoxic conditions to evaluate their sickling properties. Briefly, RBCs were resuspended in ID-CellStab stabilization solution for red cells (BIORAD) and exposed to an oxygen-deprived atmosphere for ≥ 60 min. Images were captured using the AxioObserver microscope and the Zen software (Zeiss) at a magnification of 40X. Images were analyzed with ImageJ to determine the percentage of sickling RBCs per field of acquisition in the total RBC population.

### Quantitative phase image microscopy of RBCs

At day 19 to 21 of the terminal erythroid differentiation, RBCs were collected, resuspended in PBS, and placed in a 4 well μ-Slide (ibidi). Quantitative phase images of label-free RBCs were taken using the SID4 HR GE camera (Phasics, Saint-Aubin, France) with an inverted microscope (ECLISPE Ti-E, Nikon) and a x40/0.60 objective. For each image, a segmentation procedure was performed using the BIO-Data R&D software (version 2.7.1.46) to isolate individual RBCs (Ramadier et al., 2021). For β-thalassemia samples, enucleated RBCs were selected based on their surface density (dry mass/surface), discarding nucleated cells with a surface density (pg/mm^2^) >0.376 (βAS3) and >0.4 (βAS3/miRBCL11A) for β^+^/β^+^_#1_ and >0.358 (Mock), >0.368 (βAS3), and >0.365 (βAS3/miRBCL11A) for β^0^/β^0^_#2_. Dry mass (pg), surface (mm^2^), and perimeter (mm) were measured for each enucleated RBC.

### Vector integration site Analysis

G-CSF mobilized CD34^+^ from healthy donors were thawed, cultured, and transduced as described above. After 6 days, DNA was extracted and 3′ LTR vector-genome junctions were amplified by ligation-mediated PCR (LM-PCR) and sequenced as described in Poletti et al., 2018 (Poletti et al., 2018). Comparative gene ontology analysis (GO Biological Processes BP5) was performed with DAVID tool on target genes (defined by read count >95th percentile). Data are available online in the NCBI Sequence Read Archive (SRA accession: PRJNA842958).

### RNA- and miRNA-sequencing analysis

Total RNA was extracted from erythroblasts collected at day 13 of the erythroid differentiation using the Quick-RNA MicroPrep Kit (Zymo Research). This protocol includes DNase treatment of the extracted samples. The concentration and the purity of the total RNA were measured using the Xpose spectrophotometer (Trinean).

For the RNA-seq experiments, the integrity of the RNA was evaluated by capillary electrophoresis using the Fragment Analyzer (Agilent). Before preparing the RNA-seq libraries, total RNA was treated with the Heat-labile double strand DNase (HL-dsDNase, ArcticZyme) to remove any potential residual genomic DNA contamination. Then, the Ovation Universal RNA-Seq System (Tecan) was used to prepare the RNA-seq libraries from 35 ng of total RNA following the manufacturer’s protocol. Briefly, the reverse transcription and the second strand synthesis were followed by a fragmentation step and the ligation of Illumina compatible indexed adaptor coupled to strand selection enzymatic reaction to retain the information on the orientation of the transcripts. Next, Insert Dependent Adaptor Cleavage (InDA-C) specific primers were used to deplete human ribosomal RNA transcripts before PCR enrichment. Finally, an equimolar pool of the final indexed RNA-Seq libraries was sequenced on an Illumina NovaSeq6000 (paired-end sequencing, 2×100 bp) and ~100 million paired-end reads per library were produced. Read quality was verified using FastQC (v. 0.11.9; http://www.bioinformatics.babraham.ac.uk/projects/fastqc/). Raw reads were trimmed for adapters and low-quality tails (quality < Q20) with BBDuk (v. 38.92; sourceforge.net/projects/bbmap/); moreover, the first 10 nucleotides were force-trimmed for low quality. Reads shorter than 35 bp after trimming were removed. Reads were subsequently aligned to the human reference genome (hg38) using STAR (v. 2.7.9a; Dobin et al., 2013). Raw gene counts were obtained in R-4.1.1 using the *featureCounts* function of the *Rsubread* R package (v. 2.6.4; Liao et al., 2014) and the GENCODE 38 basic gene annotation for hg38 reference genome. Transgene sequence and annotation were added to the hg38 reference genome and annotation to quantify their expression. Gene counts were normalized to counts per million mapped reads (CPM) and to fragments per kilobase of exon per million mapped reads (FPKM) using the *edgeR* R package (v. 3.34.1; Robinson et al., 2010); only genes with a CPM greater than 1 in at least 2 samples were retained for differential analysis. Differential gene expression analysis was performed using the *glmQLFTest* function of the *edgeR* R package, using donor as a blocking variable. Genes with FDR < 0.05 and absolute log2 fold change ≥ 1 were defined as differentially expressed.

For the miRNA-seq experiments, 50 ng of total RNA was used to prepare miRNA libraries using the QIAseq miRNA Library Kit (Qiagen). The amount and size of the libraries were evaluated using the TapeStation 2100 (Agilent). Libraries were mixed in equimolar ratio and sequenced on a NextSeq500 sequencing instrument. Raw reads were demultiplexed and FASTQ files were generated using the bcl2fastq software (Illumina Inc.) Read quality was evaluated using FastQC (v. 0.11.9; http://www.bioinformatics.babraham.ac.uk/projects/fastqc/). UMI extraction and adapter removal was performed using UMI-Tools (v. 1.1.2); then, reads <16 bp were removed using BBDuk (v. 38.92; sourceforge.net/projects/bbmap/). Reads were subsequently aligned to the human reference genome (hg38) using STAR (v. 2.7.9a; Dobin et al., 2013). Alignments are then de-duplicated using UMI-Tools (v. 1.1.2). UMI-based counts were obtained in R-4.1.1 using the *featureCounts* function of the *Rsubread* R package (v. 2.6.4; Liao et al., 2014) and the miRBase annotation (v22; Kozomara et al., 2019) for hg38 reference genome. miR7m sequence and annotation were added to the hg38 reference genome and annotation to quantify its expression. UMI-based counts were normalized to counts per million mapped reads (CPM) using the *edgeR* R package (v. 3.34.1; Robinson et al., 2010); only genes with a CPM >1 in at least 2 samples were retained for differential analysis. Differential gene expression analysis was performed using the *glmQLFTest* function of the *edgeR* R package, using donor as a blocking variable. Genes with FDR < 0.05 and absolute log2 fold change ≥ 1 were defined as differentially expressed.

RNA-seq and miRNA-seq data are available in the Gene Expression Omnibus repository under the accession number GSE205074.

### HSPC xenotransplantation in NBSGW mice

NOD.Cg-KitW-41JTyr +PrkdcscidIl2rgtm1Wjl/ThomJ (NBSGW) mice were housed in a pathogen-free facility. Mock- or LV-transduced non-mobilized SCD CD34^+^ cells (5×10^5^ cells per mouse) were transplanted into nonirradiated NBSGW male and female mice of 5 to 7 weeks of age via retro-orbital sinus injection. NBSGW mice were conditioned with busulfan (Sigma, St Louis, MO, USA) injected intraperitoneally (10 mg/kg body weight/day) 24 h before transplantation. Neomycin and acid water were added in the water bottle. 18 weeks after transplantation, NBSGW primary recipients were sacrificed. Cells were harvested from bone marrow, thymus and spleen and stained with antibodies against murine and human surface markers [murine CD45 (1/50 mCD45-VioBlue), Miltenyi Biotec; human CD45 (1/50 hCD45-APCvio770), Miltenyi Biotec; human CD3 (1/50 CD3-APC), Miltenyi Biotec; human CD14 (1/50 CD14-PE-Cy7), BD Biosciences; human CD15 (1/50 CD15-PE), Miltenyi Biotec; human CD19 (1/100 CD19-BV510); human CD235a (1/50 CD235a-PE), BD Biosciences] and analyzed by flow cytometry using the MACSQuant analyzer (Miltenyi Biotec) and the FlowJo software (BD Biosciences). Human bone marrow CD45+ cells were sorted by immunomagnetic selection with AutoMACS (Miltenyi Biotec) after immunostaining with the CD45 MicroBead Kit (Miltenyi Biotec). All experiments and procedures were performed in compliance with the French Ministry of Agriculture’s regulations on animal experiments and were approved by the regional Animal Care and Use Committee (APAFIS#2019061312202425_v4). Mice were housed in a temperature (20–22 °C) and humidity (40–50%)-controlled environment with 12 h/12 h light/dark cycle and fed ad libitum with a standard diet.

### Statistical analyses

Statistical analyses were performed when the total number of replicates for each group was ≥3 using GraphPad Prism version 9. We used Shapiro-Wilk test to evaluate if data follow a Gaussian distribution. For comparison between two groups or more following a Gaussian distribution, we used an unpaired t test to compare two averages or one- or two-way ANOVA test, followed by multiple comparisons, to compare three or more averages. For non-Gaussian distributed data, we used non-parametric tests, namely Mann-Whitney test for comparing two groups and Kruskal-Wallis for multiple comparison. P values are indicated with asterisks (*P<0.05, **P<0.01, ***P<0.001), and ns (not significant) reflects P>0.05.

## Results

### Development of lentiviral vectors co-expressing the βAS3 transgene and the miRBCL11A

To achieve therapeutic Hb levels for both β-thalassemia and SCD, we developed bifunctional LVs allowing *HBG* de-repression through an amiR targeting the fetal Hb (HbF) repressor *BCL11A* and the concomitant expression of the βAS3 transgene. In particular, we used an amiR (miRBCL11A) (Brendel et al., 2020, 2016; Guda et al., 2015) targeting the extra-large BCL11A isoform (BCL11A-XL) responsible for *HBG* silencing (Liu et al., 2018; Trakarnsanga et al., 2014; Zhu et al., 2012), thus avoiding the potential side effects due to the knock-down of other BCL11A isoforms. This amiR is composed of a shRNA embedded in the miR-223 backbone that has been extensively optimized to improve miRNA processing and reduce off-target binding by stringent strand selection (Amendola et al., 2009; Brendel et al., 2016; Guda et al., 2015) (**Figure 1A**). More specifically, the miR-223 backbone favors the selection of the guide strand (the strand that recognizes the target mRNA) over the passenger strand, which is instead degraded. In the case of the miRBCL11A, the guide strand of this amiR targets the 3’ end of the coding sequence of BCL11A-XL mRNA.

**Figure 1:**
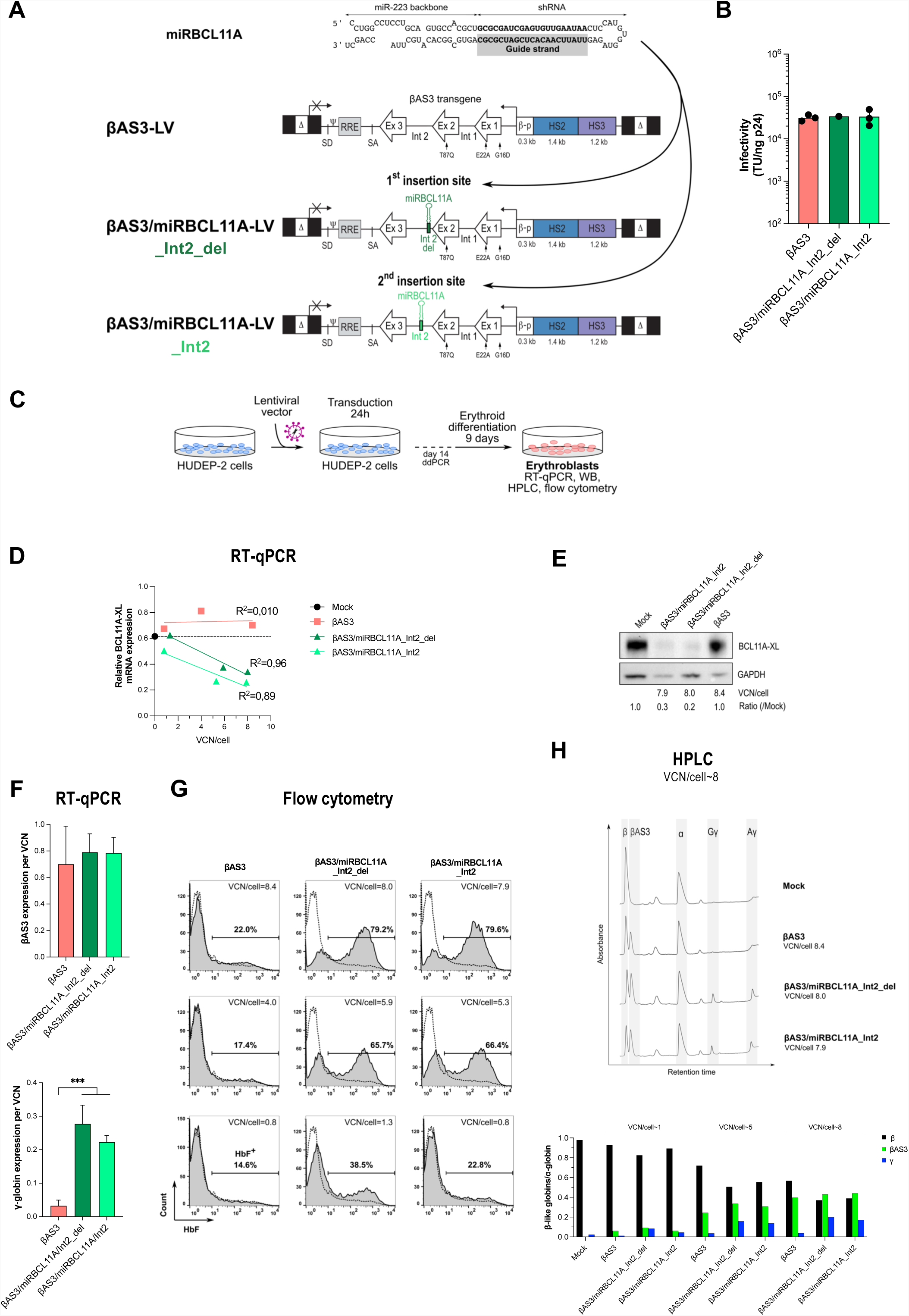
Bifunctional lentiviral vectors combining gene addition and gene silencing allow BCL11A-XL silencing, γ-globin de-repression, and βAS3 expression in HUDEP-2 cells. **(A)** The amiR (miRBCL11A) is composed of a shRNA embedded in the miR-223 backbone and targets *BCL11A-XL* mRNA after processing. The guide strand is responsible for the recognition of the target mRNA once loaded on the RISC complex. To create the novel bifunctional lentiviral vectors, we inserted miRBCL11A in two distinct sites of intron 2 of the βAS3 transgene (in βAS3-LV): Int2_del or Int2 (βAS3/miRBCL11A-LVs). Δ, deleted HIV-1 U3 region; SD and SA, HIV splicing donor and acceptor sites; Ψ, HIV-1 packaging signal; RRE, HIV-1 Rev responsive element; Ex, exons of the human *HBB*; β-p, promoter of *HBB*; HS2, 3, DNase I hypersensitive site 2, and 3 of human *HBB* LCR; arrows indicate the mutations introduced in exon 1 (generating G16D and E22A amino acid substitutions) and exon 2 (generating T87Q amino acid substitution). **(B)** Infectivity (TU/ng p24) was calculated based on infectious (TU/ml) and physical (p24 antigen ng/ml) titers (**Figure S1**) (n=1-3 independent LV productions for βAS3/miRCL11A_Int2_del, and βAS3 and βAS3/miRBCL11A_Int2, respectively). **(C)** HUDEP-2 cells were transduced at increasing MOI (1, 5, 10 and 15) for 24h. VCN/cell was measured 14 days after transduction by ddPCR. Mock- or LV-transduced cells were differentiated into mature erythroblasts for 9 days to evaluate *BCL11A-XL* and globin expression by RT-qPCR, WB, HPLC and flow cytometry. **(D-E)** *BCL11A-XL* expression measured by RT-qPCR (D) and western blot (E) in mock- and LV-transduced HUDEP-2 cells during differentiation. mRNA levels were normalized to *LMNB2* expression and protein levels to GAPDH (n=1 for Mock and n=1-3 independent biological replicates for LV-transduced cells for WB and RT-qPCR, respectively). BCL11A-XL mRNA expression was not significantly different between cells transduced with the βAS3/miRBCL11A_Int2_del- and Int2-LV (linear regression). **(F)** βAS3- (upper panel) and γ-globin (lower panel) mRNA levels were measured by RT-qPCR in LV-transduced HUDEP-2 cells after 9 days of differentiation (n=3-4 independent biological replicates for the 3 LVs). Globin mRNA levels were normalized to *HBA* expression. We plotted βAS3-globin and γ-globin mRNA levels per VCN. No significant statistical difference in βAS3 expression was observed between the 3 LVs, while γ-globin mRNA levels were significantly higher in βAS3/miRBCL11A_Int2_del- and βAS3/miRBCL11A_Int2-transduced cells than in βAS3-transduced samples). No significant statistical difference was observed between βAS3/miRBCL11A transduced samples (One-way ANOVA test; *** P<0.001 and ns, not significant). **(G)** Flow cytometry analysis of HbF expression in CD235a^high^ HUDEP-2 cells after 9 days of differentiation [mock sample: dotted line (n=1); LV-transduced samples as indicated: solid line (n=3 independent biological replicates per LV)]. **(H)** Globin expression determined by RP-HPLC in mock- (n=1) and LV-transduced cells (n=3 independent biological replicates per LV). The histogram shows β-like-/α-globin ratios. We observed an increased expression of therapeutic γ-globin and a similar expression of βAS3-globin in cells transduced with the bifunctional vectors compared to control cells (mock- or βAS3-transduced cells).

We have previously shown that amiRs can be expressed using Pol II promoters (Amendola et al., 2009). Therefore, we inserted miRBCL11A in the second 256 nucleotide-long intron of the βAS3 transgene to express it under the control of the *HBB* promoter and 2 potent enhancers derived from the *HBB* locus control region (βAS3 LV design; see Weber et al., 2018), thus reducing potential amiR toxicity by limiting its expression to the erythroid lineage (**Figure 1A**). To avoid negative effects on βAS3 RNA expression or processing (e.g., splicing and 3’-end formation), we tested only two different positions in the βAS3 intron 2 to insert the amiR (Int2_del and Int2). These regions are not known to be involved in β-globin RNA expression and splicing and far enough from the last 60 nucleotides of intron 2 known to be required for efficient 3’-end mRNA formation (Antoniou et al., 1998) (**Figure 1A**). Both physical and infectious titers of amiR-containing LVs (βAS3/miRBCL11A_Int2_del and _Int2) were similar to the original vector expressing only the βAS3 transgene (βAS3 LV) (**Figure S1**). Therefore, the insertion of the amiR did not impact viral particle production and infectivity (**Figure S1** and **Figure 1B**).

These LVs were then tested in a human erythroid progenitor cell line (HUDEP-2) (**Figure 1C**). To assess the potential impact of the amiR on gene transfer efficiency, HUDEP-2 cells were transduced at increasing multiplicities of infection (MOI) with all LVs. VCN analysis by ddPCR revealed that neither the insertion of the miR, nor its position in intron 2 affects gene transfer efficiency (**Figure S2**).

Mock- and LV-transduced HUDEP-2 cells were terminally differentiated into mature erythroblasts (**Figure 1C**). *BCL11A-XL* mRNA expression was decreased in HUDEP-2 cells transduced with LVs containing the amiR compared with control cells (mock-transduced or transduced with βAS3 LV) (**Figure 1D**). *BCL11A-XL* mRNA downregulation led to a strong decrease of BCL11A-XL at protein level in cells transduced at a high VCN/cell (**Figure 1E**). These results demonstrated that the amiR is expressed in the frame of the βAS3-expressing LVs and is able to reduce *BCL11A-XL* expression. Importantly, βAS3 mRNA was expressed at similar levels in samples transduced with amiR-containing LVs or with the control βAS3 LV, showing that neither the insertion of the amiR nor its position in βAS3 intron 2 affects transgene expression (**Figure 1F**).

To evaluate if BCL11A-XL silencing was associated with *HBG* re-activation, we measured *HBG* mRNA expression levels in terminally differentiated HUDEP-2 cells. *HBG* expression was substantially higher in mature erythroblasts transduced with amiR-expressing LVs than in cells transduced with the βAS3 LV (**Figure 1F**). Flow cytometry showed that both the percentage of HbF^+^ populations and HbF content (measured as mean fluorescence intensity, MFI) were increased in samples transduced with LVs expressing the miR targeting BCL11A compared to controls (**Figure 1G**). RP-HPLC analysis of single globin chains showed increased γ-globin levels upon BCL11A-XL silencing and equivalent βAS3 expression in all transduced samples (**Figure 1H**). Importantly, we observed a decrease in the levels of the endogenous adult β-globin (**Figure 1H**), which could be beneficial in the context of SCD to reduce the expression of the mutant β^S^-globin.

Overall, these data showed that LVs expressing a βAS3 transgene and an amiR targeting BCL11A-XL could reactivate γ-globin expression without altering βAS3 production in an adult erythroid cell line model. These bifunctional LVs can be exploited to achieve therapeutic Hb levels in both β-thalassemia and SCD. For further studies in primary patient cells and for the development of bifunctional vectors expressing alternative amiR (i.e., against *HBB*), we selected the Int2 insertion position. Although γ-globin expression was similar between the two vectors, βAS3/miRBCL11A_Int2 LV (hereafter named βAS3/miRBCL11A) tended to induce a better BCL11A-XL downregulation even at low VCN/cell, suggesting a better amiR production associated with the Int2 configuration (**Figure 1D**).

### Downregulating *BCL11A-XL* modestly increases the therapeutic potential of βAS3-expressing **LVs**

First, we transduced adult G-CSF-mobilized HSPCs derived from two β-thalassemia patients (with a β^+^/β^+^ or a β^0^/β^0^ genotype) with the bifunctional LV harboring the amiR against *BCL11A-XL* (βAS3/miRBCL11A). As control vectors, we used βAS3 LV and an LV containing a non-targeting (nt) amiR (that does not recognize any human sequence) inserted in position Int2 of βAS3 intron 2 (βAS3/miRnt). Mock- and LV-transduced HSPCs were terminally differentiated into mature RBCs (**Figure 2A**). Transduction efficiency by βAS3/miRBCL11A and control vectors was similar in erythroid liquid cultures (VCN of ~1 and ~2 in the β^+^/β^+^ and the β^0^/β^0^ patient, respectively), indicating that infectivity was not affected by the introduction of the amiR also in primary HSPCs (**Figure 2B** and **Figure S3A**). HSPCs were plated in clonogenic cultures (colony forming cell [CFC] assay) allowing the growth of erythroid (BFU-E) and granulomonocytic (CFU-GM) progenitors. Importantly, LV transduction did not alter the growth and multilineage differentiation of hematopoietic progenitors (**Figure S3B**). VCN in BFU-E and CFU-GM (~1 and ~2 in the β^+^/β^+^ and the β^0^/β^0^ patient, respectively) was similar to the values observed in the erythroid liquid culture, with ~50% of transduced progenitors (**Figure S3B** and **S3C**).

**Figure 2:**
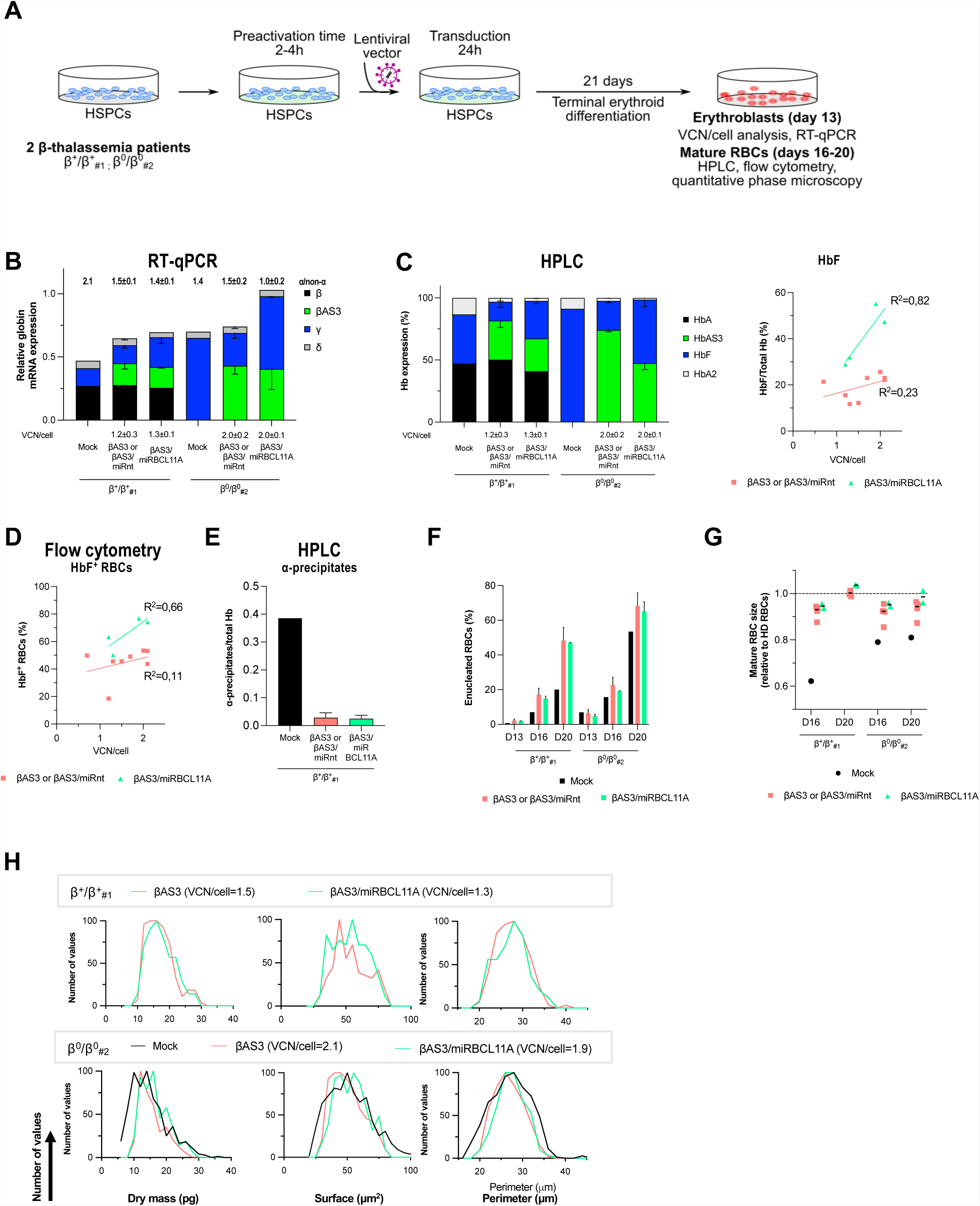
Testing a miRBCL11A-expressing bifunctional LV in β-thalassemic cells. **(A)** HSPCs from two β-thalassemia donors (β^+^/β^+^_#1_ and β^0^/β^0^_#2_) were either mock-transduced (Mock; n=1 per donor) or transduced with control (βAS3 (n=2) or βAS3/miRnt (n=2); in total n=4 independent biological replicates per donor) or bifunctional (βAS3/miRBCL11A; n=2 independent biological replicates per donor) vectors for 24 hours (n=1-4 independent biological replicates per donor). After transduction, cells were differentiated towards the erythroid lineage for 21 days. VCN/cell was measured 14 days after transduction by ddPCR in erythroblasts. Globin expression (RT-qPCR, HPLC, flow cytometry) and RBC properties (flow cytometry and quantitative phase microscopy) were evaluated in erythroblasts or mature RBCs. βAS3- and βAS3/miRnt-transduced samples were pooled in all the analyses (B-G). **(B)** Globin mRNA levels measured by RT-qPCR in mock- and LV-transduced HSPC-derived erythroblasts after 13 days of differentiation. Globin mRNA levels were normalized to *HBA*. α/non-α ratios (mean±SD) are indicated on the top of each histogram. For calculating the α/non-α ratio, globin expression was normalized to *GAPDH*. (n=1-4 independent biological replicates per donor) **(C)** Hemoglobin expression determined by CE-HPLC in RBCs after 16 days of differentiation (n=1-4 independent biological replicates per donor). Linear regression of HbF expression *vs* VCN/cell confirmed HbF reactivation in RBCs derived from βAS3/miRBCL11A-transduced HSPCs (Linear regression, *P=0.0396, n=4-8 independent biological replicates). **(D)** Proportion of HbF^+^ RBCs, quantified by flow cytometry in RBCs after 19 days of differentiation (n=4-8 independent biological replicates). **(E)** Analysis of α-globin precipitates by CE-HPLC in β^+^/β^+^ _#1_ patient RBCs. We calculated the proportion of α-globin precipitates over the total Hb tetramers (mock: n=1, βAS3 or βAS3/miRnt: n=4, and βAS3/miRBCL11A: n=2 independent biological replicates). **(F-G)** Enucleation (F) and RBC size (G) measured by flow cytometry along the differentiation (n=1-4 independent biological replicates per donor). RBC size was not analyzed in the β^+^/β^+^_#1_ mock sample because of the low number of enucleated cells. **(H)** RBC parameters [Dry mass (pg), Surface (μm^2^), and Perimeter (μm)] were extracted using the BIO-Data software from images taken with the Phasics camera after 19 days of differentiation for β^+^/β^+^_#1_ (upper panel) and β^0^/β^0^ (lower panel) samples. VCN/cell (mean±SD) is indicated in the legend. Analyses were performed on enucleated RBCs, and data were normalized to the total number of enucleated RBCs and reported as overlaid histograms. Phase microscopy analysis was not performed in the β^+^/β^+^_#1_ mock sample because of the low number of enucleated cells.

To evaluate the therapeutic potential of this strategy, we measured globin expression in mock- and LV-transduced erythroid liquid cultures. Of note, the β^0^/β^0^ samples showed high basal *HBG* levels and a less severe *in vitro* phenotype compared to the β^+^/β^+^ samples (possibly due to *in vitro* culture conditions, stress erythropoiesis or potential genetic modifiers of HbF expression present in this patient and a single 3.7-kb *HBA* gene deletion, which taken together possibly mitigate the β-thalassemic phenotype) (**Figure 2B** and see **Figure 2E-H**; Sripichai et al., 2008). However, γ-globin mRNA levels were higher in both β^+^/β^+^ or a β^0^/β^0^ cells transduced with LVs containing the amiR against *BCL11A-XL* than in control cells transduced with control LVs (**Figure 2B**). Of note, βAS3 expression was not affected in cells expressing miRBCL11A, confirming no impact of the amiR insertion on βAS3 mRNA production (**Figure 2B**). Moreover, *HBG* gene de-repression combined to βAS3 expression led to an improvement of the α/non-α globin mRNA ratio in cells transduced with the bifunctional LV compared to mock- or control LV-transduced cells (**Figure 2B**). The relative proportion of HbF, measured by CE-HPLC, was also increased as a consequence of *HBG* de-repression in cells expressing the amiR targeting *BCL11A-XL* (**Figure 2C**). Finally, flow cytometry revealed that the bifunctional LV increases the proportion of HbF^+^ RBCs (**Figure 2D**).

We then evaluated if the expression of Hb tetramers containing βAS3 (HbAS3) combined to HbF reactivation could improve β-thalassemia RBC clinical phenotypes. Alpha-precipitates detected only in cells differentiated from the β^+^/β^+^ patient were strongly decreased upon treatment with both βAS3/miRBCL11A and control vectors (**Figure 2E**). Similarly, enucleation rate and RBC size, known to be reduced in β-thalassemia were improved in samples derived from HSPCs transduced with either βAS3/miRBCL11A or the control LVs (**Figures 2F** and **2G**). No changes in the expression of erythroid surface markers along the differentiation were observed upon transduction with bifunctional LVs compared to control samples, as measured by flow cytometry (**Figure S4**). Enucleated RBC parameters evaluated by phase microscopy were similar in samples derived from HSPCs transduced with either βAS3/miRBCL11A or the control LVs (**Figure 2H**). Of note, for the β^0^/β^0^ patient, RBC parameters were improved in LV-transduced samples compared to mock-transduced cells, while untreated β^+^/β^+^ cells could not be analyzed due to the low proportion of enucleated RBCs at the end of differentiation, which was rescued by LV transduction (**Figures 2F, 2G** and **2H**). In particular, for the β^0^/β^0^ patient, dry mass was increased in RBCs derived from transduced samples reflecting an amelioration of Hb content per cell, and RBC surface and perimeter (altered because of the anisocytosis and poikilocytosis typical of β-thalassemic cells) were less variable, revealing a more homogenous cell population (**Figure 2H**).

Next, we applied this therapeutic strategy to SCD HSPCs. The expression of the anti-sickling βAS3- and γ-globins and the concomitant potential decreased expression of the adult β^S^-globin could be beneficial for SCD patients (Breda et al., 2016). SCD primary HSPCs were mock-transduced or transduced with βAS3/miRBCL11A or control LVs (βAS3 or βAS3/miRnt) at increasing MOI, and differentiated into mature RBCs (**Figure 3A**). No impairment of erythroid differentiation or RBC enucleation was observed upon transduction with the bifunctional LV compared to control samples as measured by flow cytometry (**Figure S5**). In parallel, the CFC assay showed no impairment of HSPC viability or erythroid and granulo-monocytic differentiation by the miRBCL11A (**Figure S6A**). VCN was similar for all the vectors, i.e. ~1 at a MOI of 1, up to 2-4 a MOI of 5 (up to ~2 for βAS3/miRBCL11A LV up to ~4 for control vectors), while a MOI of 25 was associated with a lower VCN (**Figure S6B**)

**Figure 3:**
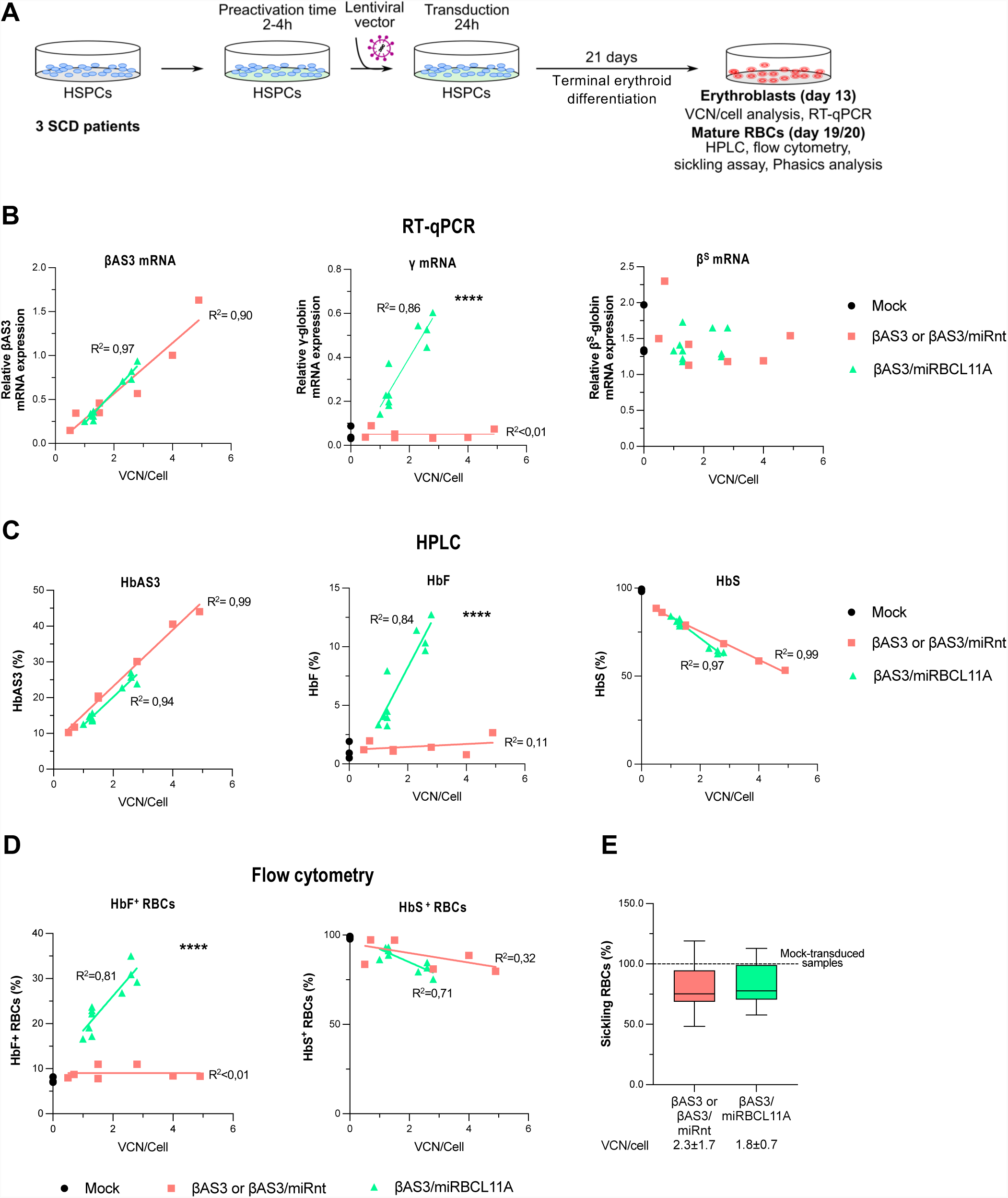
Testing a miRBCL11A-expressing bifunctional LV in SCD cells. **(A)** Mobilized (2 donors) or non-mobilized (1 donor) peripheral blood HSPCs from SCD donors were either not transduced (Mock; n=1 per donor) or transduced with control (βAS3 or βAS3/miRnt; n=2-3 independent biological replicates per donor) or bifunctional (βAS3/miRBCL11A; n=3-4 independent biological replicates per donor) vectors at different MOI for 24 hours. After transduction, cells were differentiated towards the erythroid lineage for 21 days. VCN/cell were measured 14 days after transduction by ddPCR in erythroblasts. Globin expression (RT-qPCR, HPLC and flow cytometry) and RBC differentiation markers and properties (flow cytometry and sickling assay) were evaluated in erythroblasts along the differentiation or in mature RBCs. βAS3- and βAS3/miRnt-transduced samples were pooled in all the analyses (B-F). **(B)** βAS3- (left panel), γ- (middle panel) and β^S^-globin (right panel) expression measured by RT-qPCR in erythroid precursors after 13 days of differentiation. Linear regression for γ-globin mRNA, ****P<0.0001. (n=3, 7, and 10 independent biological replicates for mock-, βAS3- or βAS3/miRnt-, and βAS3/miRBCL11A-transduced samples, respectively). **(C)** HbAS3 (left panel), HbF (middle panel) and HbS (right panel) expression measured by CE-HPLC in mature RBCs after 20 days of differentiation. Linear regression for HbF expression, ****P<0.0001. (n=3, 7, and 10 independent biological replicates for mock-, βAS3- or βAS3/miRnt-, and βAS3/miRBCL11A-transduced samples, respectively). **(D)** Proportion of HbF^+^ (left panel) and HbS^+^ (right panel) RBCs evaluated by flow cytometry. Linear regression for HbF^+^-RBCs, ****P<0.0001. (n=3, 7, and 10 independent biological replicates for mock-, βAS3- or βAS3/miRnt-, and βAS3/miRBCL11A-transduced samples, respectively). **(E)** Frequency of sickling RBCs after 1-hour-incubation at low oxygen tension (0% O_2_). VCN/cell (mean±SD) is indicated below each graph. (n=3, 7, and 10 independent biological replicates for mock-, βAS3- or βAS3/miRnt-, and βAS3/miRBCL11A-transduced samples, respectively).

RT-qPCR revealed a comparable βAS3 expression in all transduced samples and a VCN-dependent increase of γ-globin expression in cells transduced with the bifunctional LV compared to control samples (**Figure 3B**). However, no consequent β^S^-globin downregulation was induced upon *HBG* de-repression (**Figure 3B**). CE-HPLC confirmed that HbAS3 was expressed at similar levels in all transduced samples harboring similar VCN and that HbF was significantly increased in βAS3/miRBCL11A-transduced cells compared to control samples (**Figure 3C**). However, HbS expression was only modestly decreased in cells reactivating more efficiently HbF (VCN>2; **Figure 3C**). Similar results were observed in BFU-E pools (**Figure S6C**). Similarly, we observed an increased proportion of HbF^+^ RBCs upon transduction with bifunctional LV, but no significant decrease in the proportion of HbS^+^ RBCs upon HbF reactivation (**Figure 3D**). Finally, we incubated mature RBCs at low oxygen concentration to induce RBC sickling. Bifunctional and control LVs led to a comparable decrease of the frequency of sickling cells, indicating that γ-globin reactivation associated with βAS3 expression was not sufficient to further ameliorate the SCD RBC phenotype compared to βAS3 expression alone (**Figure 3E**).

In summary, we created a bifunctional LV enabling the simultaneous expression of a therapeutic transgene and an amiR down-regulating *BCL11A-XL*. βAS3/miRBCL11A LV efficiently transduces HSPCs from β-thalassemic and SCD patients and leads to HbF reactivation without impairing βAS3 transgene production. HbF reactivation combined to βAS3 expression modestly improved the β-thalassemia phenotype *in vitro*, while in SCD cells the potential therapeutic benefit of this strategy was minimal.

### miR7m efficiently downregulates β-globin expression in erythroid cell lines

The limited therapeutic benefit of the βAS3/miRBCL11A LV in SCD could be due to the persistence of high HbS levels preventing the incorporation of the therapeutic β like-globins in Hb tetramers and the correction of the sickling RBC phenotype. To reduce HbS levels and correct the SCD cell phenotype, we used our bifunctional LV design to develop a strategy aimed at expressing βAS3 and concomitantly reduce β^S^-globin levels to favor the incorporation of βAS3 into Hb tetramers.

To generate an effective miR targeting the β-globin gene, we adapted sequences from siRNAs and shRNAs targeting *HBB* or designed miR using an online tool (https://felixfadams.shinyapps.io/miRN/; Adams et al., 2017) (**Table 1** and **Figure 4A**). These sequences were inserted in the miR-223 backbone and the different amiRs targeting the β-globin (miRHBB) were inserted within the intron 2 of the βAS3 transgene (insertion site: Int2) as previously described for miRBCL11A (**Figures 1A** and **4B**). The 6 miRHBBs generated from shRNA sequences (miRHBB: miR1, miR3, miR4, miR5, miR6 and miR7) were further modified by removing 4 nucleotides at the 5’ end of the guide strand and adding a GCGC motif at the 3’ end (miRHBBm: miR1m, miR3m, miR4m, miR5m, miR6m, miR7m) (**Table 1**). These modifications were shown to further enhance the selection of the guide strand over the passenger strand in the RISC complex, thanks to an improved 3’ end thermodynamic stability of the guide/passenger strand duplex, and therefore could potentially increase β^S^-globin silencing (Guda et al., 2015). The miR11 was adapted from a siRNA designed by Dykxhoorn and colleagues; this sequence contains a CC mismatched dinucleotide at the 5’ of the guide strand and at the 3’ end of the passenger strand to increase silencing (**Table 1**) (Dykxhoorn et al., 2006).

**Figure 4:**
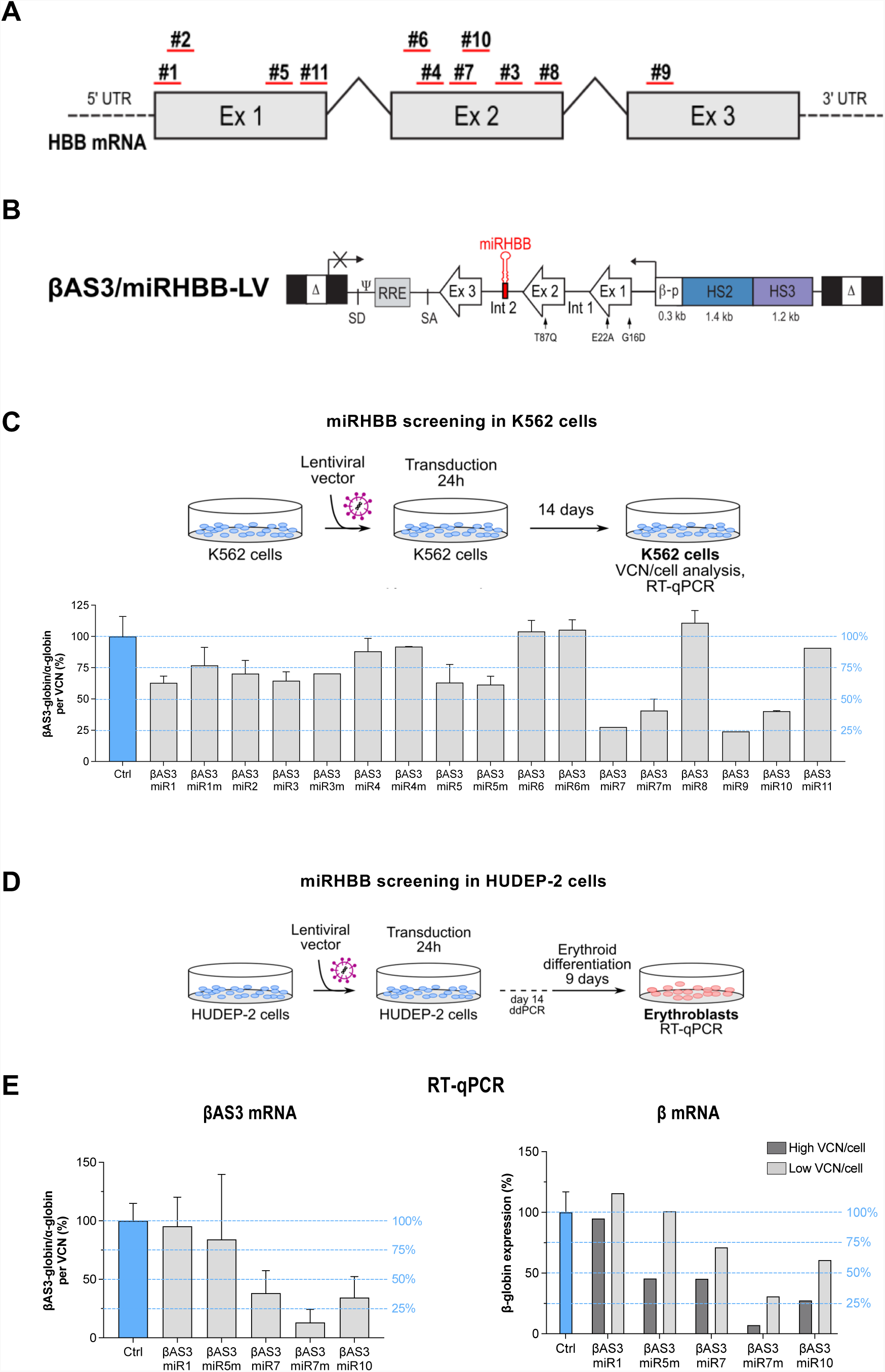
Design of a lentiviral vector expressing the βAS3-globin and an amiR targeting the β-globin. **(A)** Schematic view of the miRHBB target regions within the *HBB* mRNA. Ex1, Exon1; Ex2, Exon2; Ex3, Exon3. **(B)** Structure of the βAS3/miRHBB-LV. Δ, deleted HIV-1 U3 region; SD and SA, HIV splicing donor and acceptor sites; Ψ, HIV-1 packaging signal; RRE, HIV-1 Rev responsive element; Ex, exons of the human *HBB* gene; β-p, *HBB* promoter of; HS2, 3, DNase I hypersensitive site 2, and 3 of human *HBB* LCR; arrows indicate the mutations introduced in exon 1 (generating the G16D and E22A amino acid substitutions) and exon 2 (generating the T87Q amino acid substitution). **(C)** Screening of the 17 miRHBB in K652 cells. Cells were transduced at a MOI of 3 and 15 with control (βAS3 or βAS3/miRnt LVs, n=12 independent biological replicates) or βAS3/miRBCL11A LV (n=2 independent biological replicates, except for βAS3/miR7 and βAS3/miR9 (n=1)]. βAS3 mRNA expression was measured by RT-qPCR in K562 cells and normalized to *HBA*. We plotted the levels of βAS3 relative expression per VCN (mean±SD). **(D)** HUDEP-2 cells were transduced with control (ctrl, βAS3- or βAS3/miRnt-LV, n=4 independent biological replicates) or bifunctional (containing miR1, miR5m, miR7, miR7m, or miR10) vectors at low (2) or high (10) MOI for 24h (βAS3/miRHBB-LV, n=2 independant biological replicates per LV). VCN/cell were measured 14 days after transduction by ddPCR. Mock- or LV-transduced cells were differentiated into mature erythroblasts for 9 days to evaluate globin expression (RT-qPCR). **(E)** βAS3- (left panel) and β-globin (right panel) mRNA expression normalized to *HBA* per VCN (C, E) Results are represented as % of the control, which corresponds to the mean of the results obtained with βAS3- and βAS3/miRnt-LVs, and shown as mean±SD.

We generated a total of 17 LVs (βAS3/miRHBB) co-expressing the anti-sickling βAS3-globin and a different miRHBB. To select the most effective amiR, we transduced K562 erythroleukemic cells (that do not express endogenous β-globin) with the 17 LVs. For this amiR screening, we assessed the silencing effect of the amiR on the βAS3 transgene. The βAS3 transgene sequence was not yet modified to avoid targeting by the amiR and could be targeted by the 17 different amiRs, as there are no mismatches between the miRHBB and their target sequences within the βAS3 transgene. Therefore, we compared βAS3 expression normalized per VCN in K562 cells transduced with either βAS3/miRHBB LVs or the control LVs (βAS3 or βAS3/miRnt). K562 cells transduced with βAS3/miR7, 7m, 9 and 10

LVs showed a decrease of >50% in βAS3 mRNA expression compared to cells transduced with control LVs (**Figure 4C**). However, the cells transduced with βAS3/miR9 LV have a very low VCN/cell (<0.5) in comparison with other LVs, which could reflect a low gene transfer efficiency that hampers its use as a therapeutic vector for SCD. In cells transduced with LV βAS3/miR1 and 5m, we observed a reduction in βAS3 expression of ~30% compared to control cells (**Figure 4C**). Overall, modification of the miR sequences (Guda et al., 2015) did not increase gene silencing in K562 cells (**Figure 4C**). Based on these results we select LVs expressing miR1, miR5m, miR7, miR7m or miR10 for further experiments.

To confirm that the selected amiRs downregulate the endogenous β-globin gene, we transduced HUDEP-2, an adult erythroid progenitor cell line. These cells can be differentiated into mature erythroid precursors expressing the endogenous α and β-globin chains at both mRNA and protein levels. HUDEP-2 cells were transduced with control (βAS3 and βAS3/miRnt) and miRHBB LVs at a high (10) or a low (2) MOI and differentiated to evaluate miR efficiency in down-regulating *HBB* and βAS3 expression (**Figure 4D**). We observed a strong reduction of the βAS3 transcripts per VCN (from ~ 60% to 85%) in cells transduced with the LVs expressing miR7, miR7m and miR10, confirming the results obtained in K562 cells (**Figures 4C and 4E**). The decrease in the endogenous β-globin expression was correlated with the VCN/cell. For miR7, miR7m and miR10, we observed a reduction of the β-globin expression ranging from 60% to 85% and from 30% to 70% at a high and low VCN/cell, respectively (**Figure 4E**). On the contrary, miR1 and miR5m showed no or little activity on βAS3 and endogenous β-globin (**Figure 4E**). Of note, in HUDEP-2, the modified version of miR7 (miR7m) outperformed miR7 in terms of β-like globin silencing.

### The βAS3m/miR7m LV efficiently reduces HbS levels and corrects the sickling phenotype in RBCs differentiated from SCD primary HSPCs

Based on our results, we selected miR7m as our best performing miR in terms of β-globin silencing. Given the high sequence similarity between the β^S^-globin and the transgene, we modified the target sequence in βAS3 to avoid its silencing by the miR. To this aim, we introduced silent mutations in the βAS3 transgene sequence of βAS3/miR7m and βAS3/miRnt LV, choosing, when possible, codons commonly present in the *HBB* coding sequence (βAS3m, modified βAS3 transgene; **Figure 5A**). Overall, we introduced in the miR7m target region, a number of mutations that reduces by 33% the complementarity between the miR and the βAS3 transcript (**Figure 5A**). Notably, vector titers and infectivity were comparable between βAS3m/miR7m and βAS3m/miRnt LVs and the original βAS3 LVs, demonstrating that transgene modification does not impact the viral titer (**Figures 5B** and **S7**).

**Figure 5:**
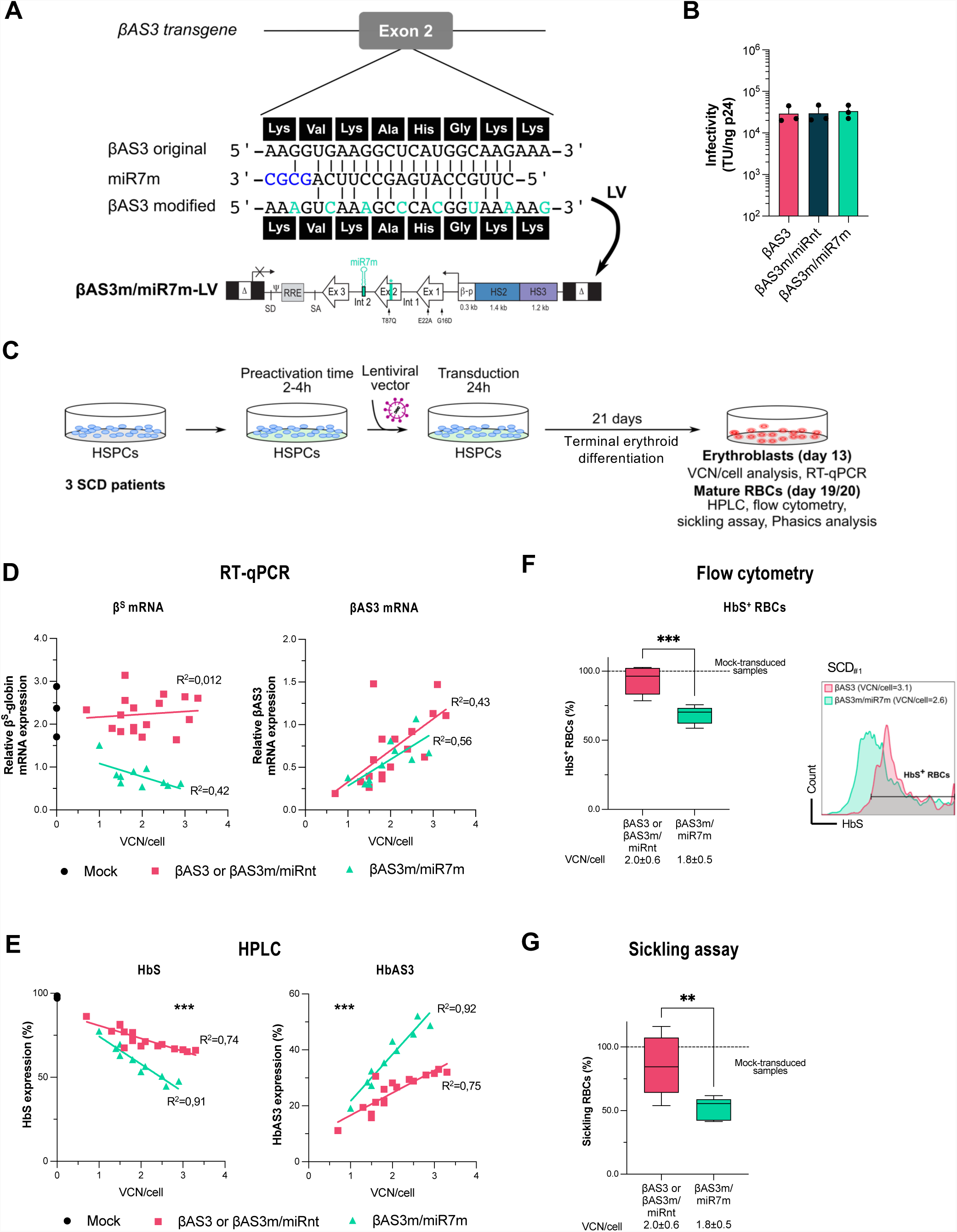
Testing the miR7m-expressing bifunctional LV in SCD HSPCs. **(A)** Structure of the βAS3m/miR7m LV. The sequence of the miR7m aligned to the βAS3 transgene is shown in the top panel. The βAS3 transgene (βAS3 original) was modified to avoid its targeting by miR7m, thus generating the βAS3 modified transgene (βAS3m) harboring mismatches (in green) with the βAS3 original transgene corresponding to silent mutations, as shown by the amino acid sequence (black boxes). Δ, deleted HIV-1 U3 region; SD and SA, HIV splicing donor and acceptor sites; Ψ, HIV-1 packaging signal; RRE, HIV-1 Rev responsive element; Ex, exons of the human *HBB*; β-p, *HBB* promoter; HS2, 3, DNase I hypersensitive site 2, and 3 of human *HBB* LCR; arrows indicate the mutations introduced in exon 1 (generating the G16D and E22A amino acid substitutions) and exon 2 (generating the T87Q amino acid substitution). **(B)** Infectivity (TU/ng p24) was calculated based on infectious (TU/ml) and physical (p24 antigen ng/ml) titers (**Figure S7**). Data are shown as mean±SD. (n=3 independent LV productions per LV) **(C)** Mobilized (2 donors) or non-mobilized (1 donor) peripheral blood HSPCs from SCD donors were either mock-transduced (Mock; n=1 per donor) or transduced with βAS3 or βAS3m/miRnt (n=5-6 independent biological replicates per donor) or βAS3m/miR7m (n=2-5 independent biological replicates per donor) LV at different MOI for 24 hours. After transduction, cells were differentiated in liquid cultures towards the erythroid lineage. VCN/cell were measured 14 days after transduction by ddPCR in erythroblasts. Globin expression (RT-qPCR, HPLC and flow cytometry) and RBC differentiation markers and properties (sickling assay) were evaluated in erythroblasts along the differentiation or in mature RBCs. βAS3- and βAS3m/miRnt-transduced samples were pooled in all the analyses (D-G). VCN/cell is indicated below each graph. (D, E) 3 SCD donors (F, G) 2 SCD donors. **(D)** β^S^- and βAS3-globin mRNA expression normalized to *HBA*. (n=3, 17, and 10 independent biological replicates for mock-, βAS3- or βAS3/miRnt-, and βAS3/miRBCL11A-transduced samples, respectively). **(E)** HbAS3 and HbS expression in mature RBCs measured by CE-HPLC. Linear regression, ***P=0.0005. (n=3, 17, and 10 independent biological replicates for mock-, βAS3- or βAS3/miRnt-, and βAS3/miRBCL11A-transduced samples, respectively) **(F)** Proportion of HbS^+^-RBCs measured by flow cytometry analysis using an antibody recognizing specifically HbS. Mann-Whitney test, ***P<0.001. (n=2, 11, and 5 independent biological replicates for mock-, βAS3- or βAS3/miRnt-, and βAS3/miRBCL11A-transduced samples, respectively) **(E)** Frequency of sickling RBCs after 1-hour incubation at low oxygen tension (0% O_2_). Unpaired t-test, **P<0.01. (n=2, 11, and 5 independent biological replicates for mock-, βAS3- or βAS3/miRnt-, and βAS3/miRBCL11A-transduced samples, respectively).

We transduced primary adult HSPCs derived from SCD donors with βAS3m/miR7m- and control-LVs (βAS3m/miRnt, and βAS3) at increasing MOI. Mock- and transduced HSPCs were subjected to a CFC assay or differentiated in mature RBCs (**Figure 5C**). The number and the proportion of BFU-E and CFU-GM was similar amongst the different samples (**Figure S8A**), indicating no impairment in erythroid and granulomonocytic cell growth and differentiation. VCN/cell in BFU-E and in erythroblasts was ~2 in all the samples even at a low MOI (1/2) (**Figure S8B**). Furthermore, we determined VCN/cell in single BFU-E and showed a similar gene transfer efficiency in βAS3m/miR7m- and control LV-transduced samples (~50% at a MOI of 1/2 and ~65% at a MOI of 15; **Figure S8C**). Moreover, βAS3m/miR7m- and control LV-transduced samples displayed a similar VCN/cell distribution with the majority of BFU-E colonies harboring 1-2 VCN/cell (**Figure S8C**).

To evaluate the reversion of the SCD cell phenotype, SCD HSPCs were terminally differentiated into mature enucleated RBCs (**Figure 5C**). Efficient HSPC transduction by βAS3m/miR7m LV led to a VCN-dependent decrease of β^S^-globin transcripts in HSPC-derived erythroid cells compared to cells transduced with control LVs (**Figure 5D**). Notably, the miR specifically down-regulated β^S^-globin, without affecting βAS3 expression (**Figure 5D**). CE-HPLC analysis showed that β^S^-globin gene downregulation led to a significant decrease of HbS that was associated with a substantial increase of the therapeutic Hb (**Figure 5E**). These results were confirmed in BFU-E pools (**Figure S9A**). We also measured β-like globin expression in individual BFU-E derived from HD HSPCs transduced with the βAS3m/miR7m- or control-LVs. Importantly, the proportion of βAS3-globin transcripts was higher in BFU-E transduced with the βAS3m/miR7m-LV than in BFU-E transduced with the control vector at both low or high VCN/cell (**Figure S9B**).

In liquid erythroid SCD cell cultures, we observed a significant reduction of the proportion of HbS^+^ cells in βAS3m/miR7m-compared to control LV-transduced samples as measured by flow cytometry (**Figure 5F**). The increased incorporation of βAS3 in Hb tetramers and the decrease in β^S^-globin led to a better correction of the sickling phenotype in mature RBCs derived from HSPCs transduced with βAS3m/miR7m LV *vs* control LVs (**Figure 5G**). Of note, the frequency of sickling cells obtained using our bifunctional LV approaches that observed in asymptomatic SCD carriers (Ribeil et al., 2017).

Importantly, RBC enucleation and erythroid differentiation were not affected by β^S^-globin down-regulation (**Figure 6A, 6B** and **6C**). Similarly, RBC parameters, such as dry mass, surface, and perimeter evaluated by phase microscopy, showed no major change induced by β^S^ downregulation (**Figure 6D**). We only observed a minor reduction in dry mass and surface in a small fraction of RBCs (13.7%±3.8% and 8.1%±6.2% for dry mass and surface, respectively) in samples obtained from HSPCs transduced with the bifunctional βAS3m/miR7m LV (**Figure 6D**). To evaluate if the reduced dry mass could reflect a β-like-thalassemia phenotype, we measured by RP-HPLC globin chain expression and evaluated the α/non-α globin ratio typically altered in β-thalassemia (*HBB* KO sample; **Figure 6E**). Importantly, RBCs derived from HSPCs transduced with the βAS3m/miR7m or control LV showed a normal α/non-α globin ratio compared to β-thalassemic cells (**Figure 6E**).

**Figure 6:**
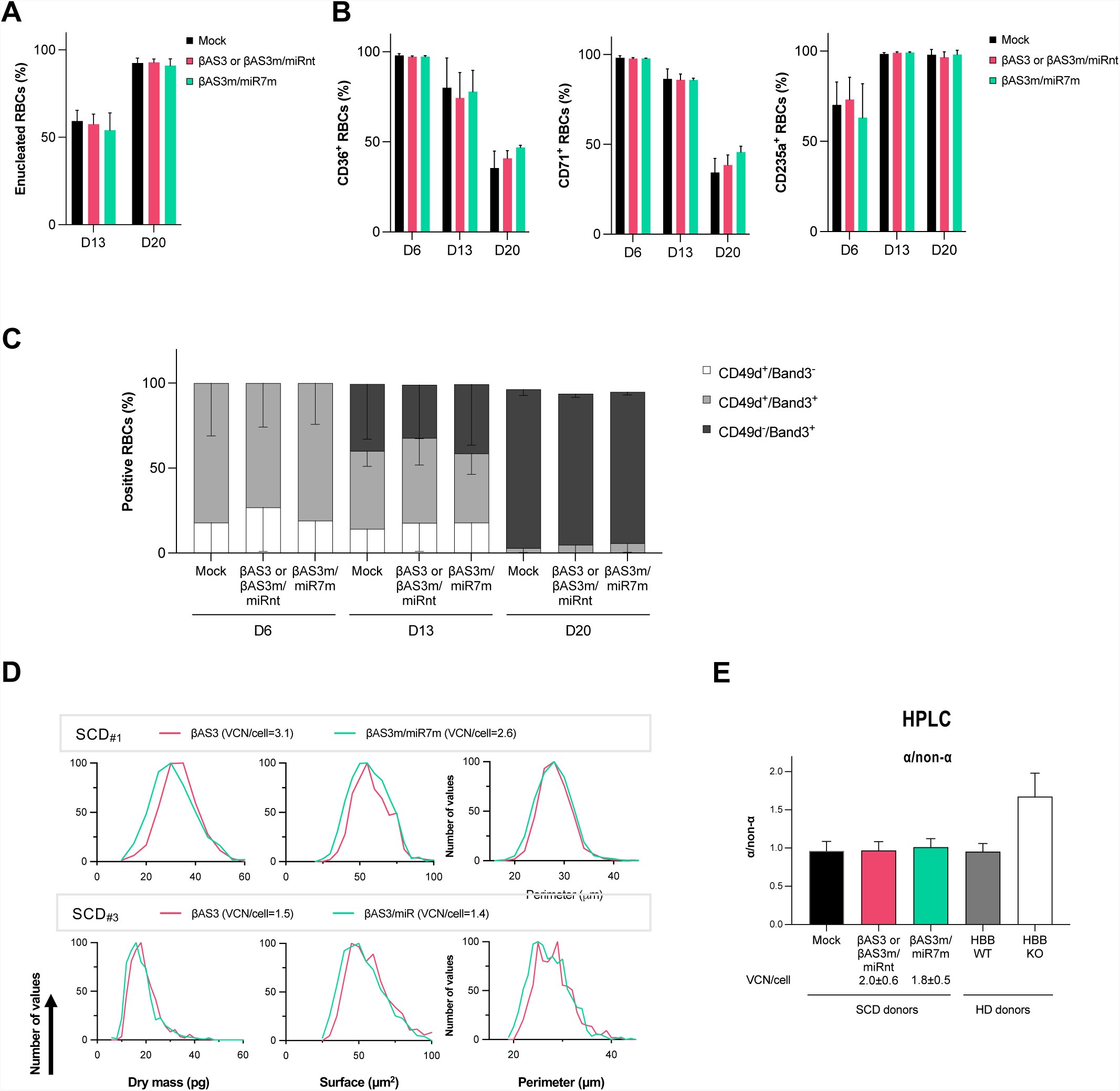
Analysis of the βAS3m/miR7m on erythroblast and RBC properties derived from SCD HSPCs. **(A)** Frequency of enucleated RBCs measured by flow cytometry at day 13 and 20 of erythroid differentiation. **(B)** Frequencies of (left) CD36^+^, (middle) CD71^+^, and (right) CD235a^+^ erythroid cells measured by flow cytometry along the differentiation. **(C)** Frequencies of CD49^+^, Band3^+^ and CD49^+^Band3^+^ cells among the CD235a^+^ RBCs measured by flow cytometry along the differentiation. **(D)** RBC parameters [Dry mass (pg), Surface (μm^2^), Perimeter (μm)] were extracted using the BIO-Data software from images taken with the Phasics camera after 19-21 days of differentiation for two SCD samples, SCD_#1_ (upper panel) and SCD_#3_ (lower panel), at a representative VCN/cell. Data were normalized to the total number of mature RBCs and reported as overlaid histograms. (**E**) α/non-α ratios (mean±SD) calculated by measuring globin expression by RP-HPLC. VCN/cell is indicated below the graph. HD samples (HD donors) are HD HSPCs that were either mock-transfected (HBB WT) or modified using a CRISPR/Cas9-gRNA RNP complex disrupting the *HBB* gene (HBB KO) (Ramadier et al., 2021). Cells were differentiated toward the erythroid lineage. βAS3- and βAS3m/miRnt-transduced samples were pooled in all the analyses (A-C, E). (A-C, E) 2 SCD donors, βAS3- and βAS3m/miRnt-LVs: VCN/cell=2.0±0.6; βAS3m/miR7m-LV: VCN/cell=1.8±0.5. (n=2, 11, and 5 independent biological replicates for mock-, βAS3- or βAS3m/miRnt-, and βAS3m/miR7m-transduced samples, respectively). (E) HD (n=2 independent biological replicates).

### The βAS3m/miR7m LV has safe integration and transcriptomic profiles in primary cells and does not affect engraftment and multi-lineage differentiation capability of HSPCs in immunodeficient mice

First, we mapped the integration sites of amiR-containing LV by LM-PCR and NGS sequencing in HD HSPCs. Our analyses revealed that βAS3m/miR7m LV showed a standard, safe lentiviral integration profile, which was essentially equivalent to that of the control βAS3 LV in terms of genome-wide distribution and top targeted genes (**Figures 7A** and **7B**). Gene ontology analysis confirmed that genes targeted by the bifunctional and control LVs have highly similar biological functions (**Figure S10**). We also performed RNA-seq and miRNA-seq on treated and non-treated erythroblasts differentiated from HSPCs derived from two SCD donors to confirm the safety of our approach and quantify the expression of the transgene and the miR7m (**Figure 7**). Transduction with the βAS3-LV modestly affected the erythroid transcriptome (89 differentially expressed genes representing 0.6% of total genes compared to mock-transduced cells; **Figure 7C**). The addition of the miRnt did not further affect gene expression (no deregulated genes between βAS3- and βAS3m/miRnt-LV transduced samples; **Figure 7C**). Only < 0.1% of total genes were modestly deregulated upon transduction with the βAS3m/miRHBB-LV compared to samples transduced with the βAS3m/miRnt-LV (**Figure 7C**). As expected, endogenous *HBB* gene expression was significantly down-regulated upon transduction with the βAS3m/miRHBB-LV (2.2-fold decrease, FDR < 0.01; **Figure 7D**). Moreover, RNA-seq experiments confirmed that βAS3 expression in the transduced samples (βAS3 or βAS3m/miR7m) was not affected by the miR7m (βAS3m/miR7m *vs* βAS3) (**Figure 7D**). As expected, the pri-miR7m was detected only in βAS3m/miR7m-transduced samples (**Figure 7D**). Similarly, the miRNAome was not affected in samples transduced with the βAS3m/miRHBB-LV compared to control samples transduced with βAS3- and βAS3m/miRnt-LV (**Figure 7E**). As expected, the mature miR7m was detected only in βAS3m/miR7m-transduced cells and its expression was in the range of endogenous miRs involved in erythroid differentiation (e.g., miRs upregulated along erythropoiesis such as miR15b-5p, miR-96-5p, miR-22-3p, or miRs downregulated along erythropoiesis, such as miR-223-5p, miR-221-3p, miR-222-3p, miR-181a-3p) (**Figure 7F**) (Papasavva et al., 2021).

**Figure 7:**
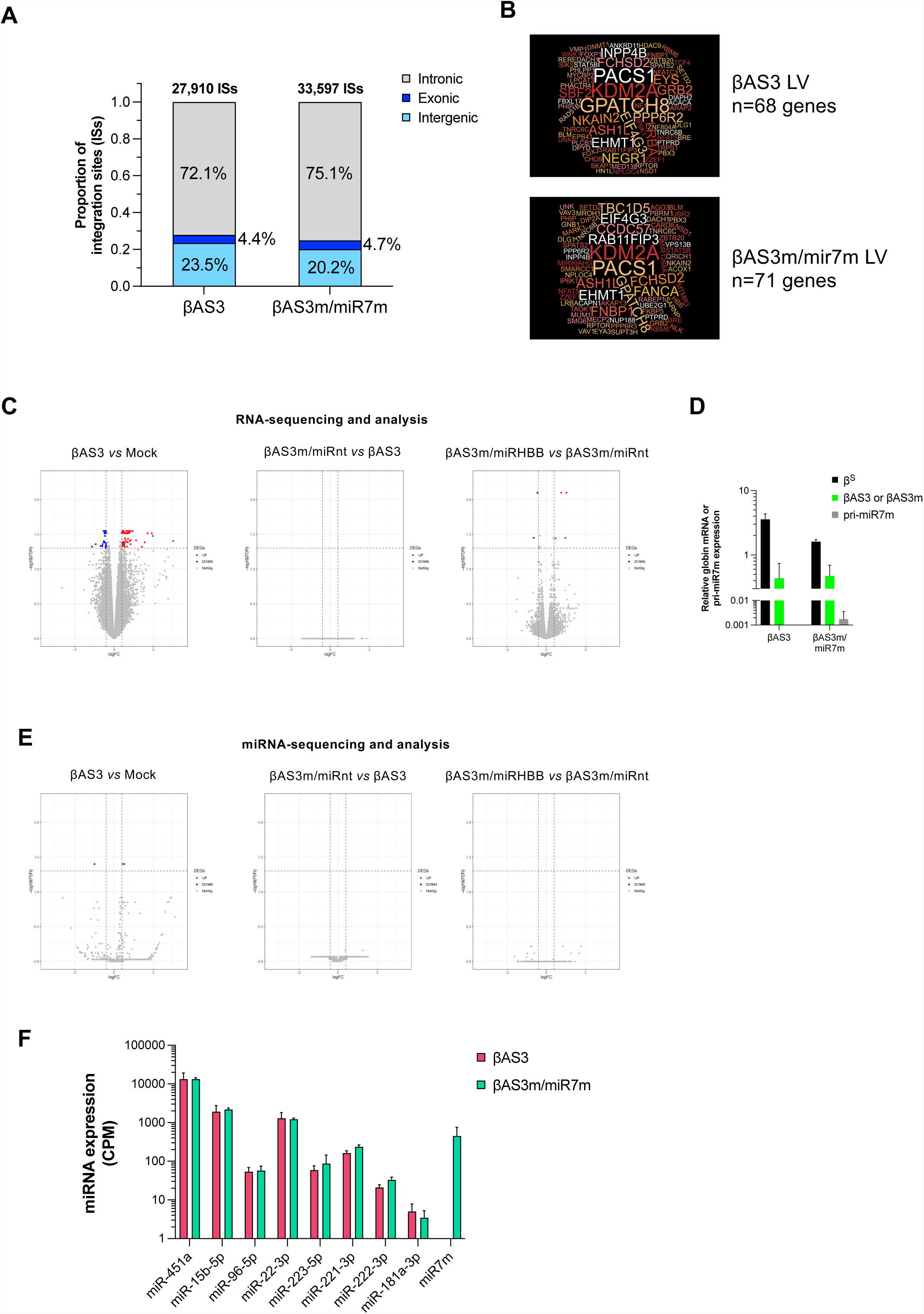
βAS3m/miR7m and βAS3 LVs displayed comparable integration and mRNA/ miRNA expression profiles. **(A)** Distribution of βAS3-LV or βAS3m/miR7m-LV integration sites in exonic, intronic and intergenic genomic regions in healthy donor HSPCs (n=1). **(B)** Top targeted genes in βAS3-LV and βAS3m/miR7m-LV-transduced HSPCs (n=1 per LV). βAS3-LV integration sites were retrieved from Poletti et al., 2018. **(C-E)** mRNA- (**C-D**) or miRNA-seq-based (**E**) analysis comparing mRNA or miRNA expression between two sample groups: βAS3 and Mock (left), βAS3m/miRnt and βAS3 (middle), βAS3m/miR7m and βAS3m/miRnt (right) (2 SCD donors were used; n=2 and 3 independent biological replicates for Mock and LV-transduced samples, respectively). Differentially expressed genes or miRs (DEGs) with an FDR < 0.05 and an absolute log2 fold-change (logFC) ≥ 1 and are highlighted in red (up-regulated, UP) or in blue (down-regulated, DOWN). Genes or miRs that are not differentially expressed are represented in grey (NotSig). (**D**) Globin (*HBB* and βAS3) mRNA and pri-miR7m expression measured by RNA-seq and normalized to *HBA* in βAS3- and βAS3m/miR7m-LV transduced samples (n=3 independent biological replicates per LV). (**F**) Expression of selected endogenous miRs involved in erythroid differentiation, which are upregulated (e.g., miR-451a, miR15b-5p, miR-96-5p, and miR-22-3p) or downregulated (e.g., miR-223-5p, miR-221-3p, miR-222-3p, and miR-181a-3p) along differentiation (Papasavva et al., 2021), and the artificial miR7m in βAS3- and βAS3m/miR7m-transduced cells measured by miRNA-seq and expressed in CPM (count per million reads) (n=3 independent biological replicates per LV).

Finally, we xenotransplantated human HSPCs from a SCD donor (either mock-transduced or transduced with the βAS3- or the βAS3m/miR7m-LV) in NBSGW mice (**Figure 8A**). The proportion of human CD45^+^ cells in the bone marrow, the spleen and the thymus was similar between the three groups, confirming that the βAS3m/miR7m-LV does not impair long-term engraftment capability of primary HSPCs (**Figure 8B**). Furthermore, engrafted βAS3m/miR7m-transpduced HSPCs were able to generate all the hematopoietic lineages (**Figures 8C-E**). The VCN/cell in the bone marrow of mice from receiving βAS3m/miR7m- or βAS3-transduced cells was not significantly different (1.5 to 1.7 for βAS3m/miR7m and 0.2 to 6.5 for βAS3; **Figure 8F**). Overall, these results validate the therapeutic potential and the safety of this novel bifunctional LV.

**Figure 8:**
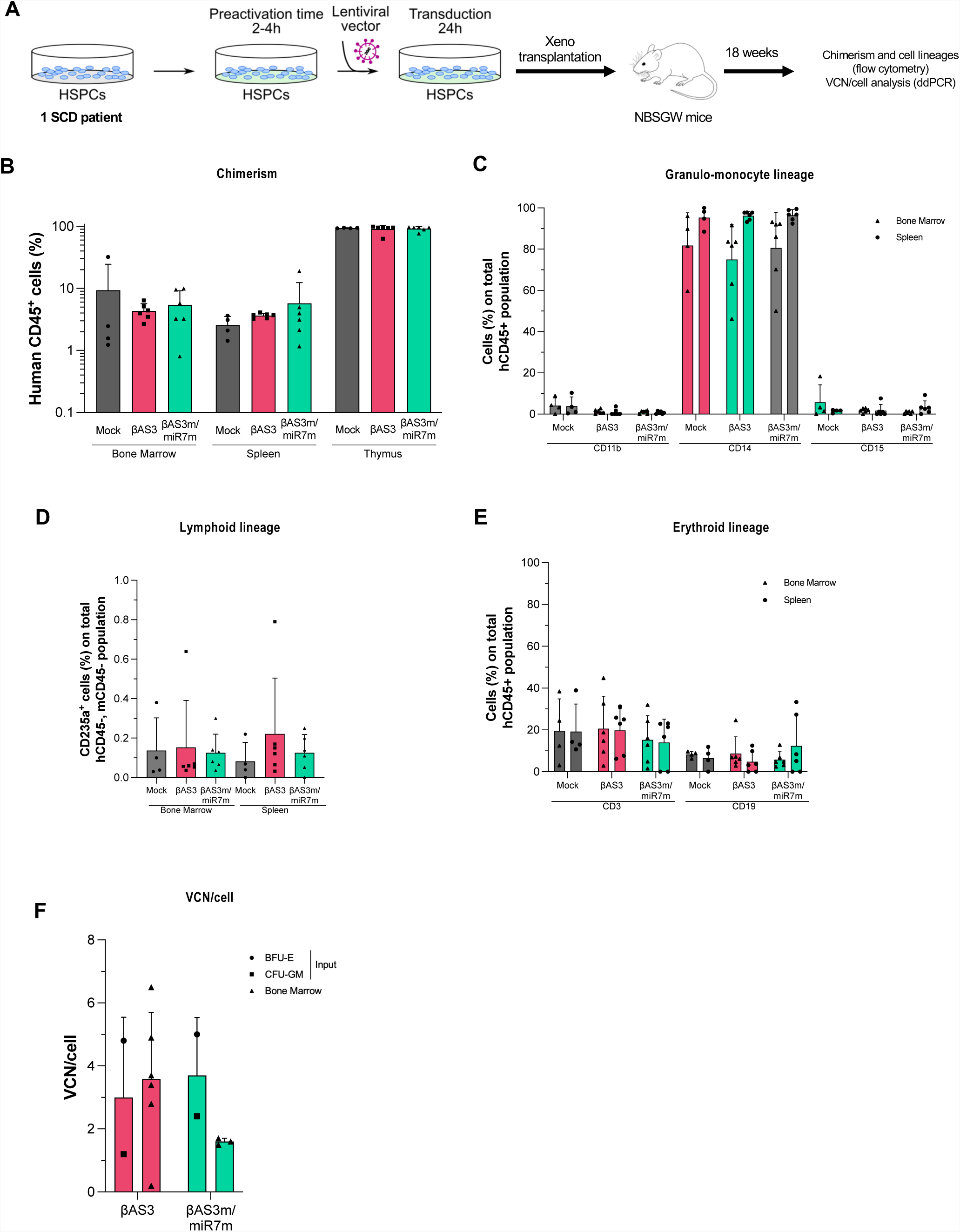
βAS3m/miR7m LV-transduced HSPCs engraft and differentiate in NBSGW mice. **(A)** Human SCD HSPCs were either mock-transduced (Mock) or transduced with βAS3 or βAS3m/miR7m LVs at a MOI of 125 for 24 hours. After transduction, cells were transplanted in NBSGW mice. After 18 weeks, chimerism, cell lineage reconstitution and VCN/cell were evaluated. **(B)** Frequency (%) of human CD45^+^ (hCD45^+^) cells in the bone marrow, the spleen and the thymus of the transplanted mice (n=4-6 mice per group). **(C)** Frequencies (%) of CD11b, CD14 and CD15 positive cells in the hCD45^+^ population in the bone marrow and in the spleen of the transplanted mice (n=4-6 mice per group). **(D)** Frequencies (%) of CD3 and CD19 positive cells in the hCD45^+^ population in the bone marrow and in the spleen of the transplanted mice (n=4-6). **(E)** Frequencies (%) of CD235a positive cells in the negative human (h) and mouse (m) CD45 population in the bone marrow and the spleen of the transplanted mice (n=4-6 mice per group). (B-E) Kruskal-Wallis test, ns. **(F)** VCN/cell measured by ddPCR in the input cells (bulk of BFU-E and CFU-GM colonies) and in the bone marrow of transplanted mice (n=3-6 mice per group). No significant statistical difference in VCN/cell was observed between the two LVs in BM cells (Unpaired t-test; ns).

## Discussion

Ongoing LV-based clinical trials for SCD revealed an amelioration of clinical phenotype associated with expression of therapeutic β-like globins via gene addition or gene silencing; however, Hb level remains lower than the normal 12-18 g/dl range in the majority of SCD and β0/β0 thalassemic patients (Esrick et al., 2021; Kanter et al., 2021; Locatelli et al., 2021; Magrin et al., 2022). In this study, we developed bifunctional LVs combining gene addition and gene silencing to improve gene therapy for β-thalassemia and SCD. The insertion of an amiR – targeting BCL11A-XL or the β^S^-globin – in the second intron of the βAS3 transgene did not affect LV titer, gene transfer efficiency and βAS3 expression in primary CD34^+^ HSPCs derived from β-hemoglobinopathy patients.

Combining βAS3 and BCL11A-XL down-regulation reactivated γ-globin but improved only the β-thalassemia phenotype compared to the mere gene addition strategy. At similar VCN, γ-globin levels were modest compared to those obtained by only expressing the shmiR targeting BCL11A-XL under the control of similar regulatory elements (Brendel et al., 2020; Esrick et al., 2021), suggesting that the processing of the intron-embedded amiR is less efficient or that βAS3 globin competes with γ-globin for the incorporation in the Hb tetramers – at least in SCD cells. Interestingly, our results are in line with a recent study describing a similar LV expressing simultaneously the T87Q β-globin (carrying only one anti-sickling amino acid) and an amiR targeting BCL11A-XL (Pires Lourenco et al., 2021). Of note, in the case of β-thalassemia, this strategy could be combined with the introduction of a second amiR targeting the α-globin, which was recently shown to improve the α/β ratio in erythrocytes derived from β-thalassemia HSPCs (Nualkaew et al., 2021). Another miR-based gene therapy strategy has recently been developed to further boost HbF expression by expressing two amiRs targeting both BCL11A or ZNF410 γ-globin repressors (Liu et al., 2022). However, the increase in HbF was limited and this therapeutic strategy modestly improved the pathological phenotype in SCD or β-thalassemic erythroid cells compared to the individual approaches.

All these therapeutic strategies are promising for β-hemoglobinopathies; however, their beneficial effect might be limited in SCD because of the competition between the therapeutic globins and the β^S^-globin for the formation of Hb tetramers. Hence, to reduce HbS levels – still high after γ-globin reactivation and/or βAS gene addition – we replaced the miRBCL11A with a miR targeting the β^S^-globin. Among the different miRs, the miR1/1m and miR2 were adapted from shRNA (Samakoglu et al., 2006) and siRNA (Dykxhoorn et al., 2006), respectively, which efficiently silenced β-globin expression. The lower expression of the miR compared to high expression rates of shRNA/siRNA could explain in part the lower silencing efficacy of these miRs. Moreover, the integration of shRNA or siRNA sequences in miR backbone could affect miR processing and therefore silencing efficiency. Nevertheless, others miRs, such as miR7m and miR10, adapted from shRNAs efficiently silenced β^S^ expression. Overall, modification of the miR sequences described by Guda and colleagues (Guda et al., 2015) did not increase βAS3 gene silencing in K562 cells, but in HUDEP-2 miR7m outperformed miR7 in reducing βAS3 and wt β-globin expression, suggesting that this design can be explored to increase miR processing and ameliorate silencing of efficient amiR.

Next, we developed the βAS3m/miR7m LV containing the most efficient miR. β^S^-globin downregulation induced by miR7m led to a reduced expression of HbS and a significant decrease of HbS^+^ RBCs. Interestingly, despite the modest BCL11A gene down-regulation using our vector design, β-globin gene silencing was highly efficient and finely tuned to avoid excessive β-globin down-regulation. Importantly, neither miR7m nor the introduction of silent mutations in exon 2 affected βAS3 expression. It has been reported in other systems that, silent mutations could alter mRNA structure and half-life or impair protein production because of the low abundance of the cognate tRNA (Kabra et al., 2022). On the contrary, elevated HbAS3 levels led to a better correction of the sickling phenotype compared to control cells. Sickle β-globin down-regulation was compensated by βAS3 expression, therefore avoiding the generation of a β-thalassemic phenotype. A modest reduction in Hb content (i.e., dry mass) was observed in a small proportion of βAS3m/miR7m-treated RBCs, but the lower Hb concentration could further reduce its polymerization and improve the RBC phenotype (Das et al., 2015; Parrow et al., 2021; Rao et al., 1983). Moreover, beneficial effects of this strategy will be more pronounced *in vivo,* owing to the positive selection of corrected RBCs likely due to their prolonged lifespan compared to non-modified SCD RBCs (Magrin et al., 2022).

It is noteworthy that here we improved LVs to boost therapeutic β-like globin levels without increasing the mutagenic vector load in HSPCs. Increasing vector load could favor therapeutic Hb formation and ameliorate the outcome of the current LV-based clinical trials; however, it potentially increases the genotoxicity risk associated with elevated LV integration sites per cell. This is particularly critical in SCD patients, who are characterized by a high mutational burden and inflammation that can be associated with clonal hematopoiesis. Recent studies reported that SCD patients have an increased risk of developing myeloid malignancies compared to the general population (Brunson et al., 2017; Seminog et al., 2016). Furthermore, three myelodysplasias associated with pro-oncogenic mutations and genomic rearrangements were observed in an ongoing gene therapy trial for SCD (Hsieh et al., 2020; Kaiser, 2021). Even if these latter events were not associated with LV integration, it is desirable to minimize the VCN/cell. Hence, our strategy, based on two therapeutic solutions to further boost therapeutic Hb, can reduce the VCN necessary to correct the SCD phenotype and thus LV-associated potential genotoxicity. Moreover, the lower VCN required to correct SCD phenotype will also diminish the cost of LV production, making this strategy potentially widely available (Coquerelle et al., 2019).

Importantly, from a safety point of view, the insertion of the amiR showed no impairment in HSPC viability and multi-lineage differentiation. βAS3m/miR7m LV showed a standard lentiviral integration profile, did not alter the transcriptome or the miRNAome of transduced erythroblasts and did not impair HSPC long-term engraftment and hematopoietic lineage reconstitution (**Figure 8**). The VCN/cell in human bone marrow cells tended to be lower in cells transduced with the βAS3m/miR7m LV compared to control cells treated with the βAS3 vector, but still remains therapeutically relevant (mean=1.6; **Figure 8F**), considering the better performance of the bifunctional LV (**Figure 5**). The safety of our therapeutic strategy is also supported by the ongoing clinical trial using LV expressing a shmiR that showed a stable gene marking over time without any adverse events associated with the drug product (Esrick et al., 2021). Notably, a similar strategy was used by Samakoglu and colleagues by co-expressing γ-globin and an shRNA (inserted in the second intron of the γ-globin transgene) targeting the β^S^-globin (Samakoglu et al., 2006). However, γ-globin does not contain an aspartic acid in position 16 that increase affinity for the a-globin chain (and is indeed likely outcompeted by βAS3). Furthermore, the pri-miRNA needs to be processed to generate a mature, functional amiR, while the shRNA is more “ready-to-be-used”. In particular, the pri-miRNA does not saturate a specific step of the endogenous miRNA pathway (Castanotto et al., 2007; Grimm, 2011). On the contrary, shRNA can cause toxicity associated with the oversaturation of cellular small RNA pathways (namely RISC incorporation), which could lead to alteration in the production of endogenous miRs. Importantly, in our study using an amiR expressed through a lentiviral vector, we did not see relevant changes in the transcriptome and miRNAome. Finally, shRNA can trigger the innate immune response and cytotoxicity (Boudreau et al., 2009; Course et al., 2020; McBride et al., 2008) while amiR are known to be associated with a lower toxicity (Borel et al., 2011; Boudreau et al., 2009; Grimm, 2011; Gröβl et al., 2014; McBride et al., 2008). Of note, the interferon-responsive gene *IRF*/9*ISGF3G* was significantly upregulated in cell treated with the shRNA (Samakoglu et al., 2006). On the contrary, we did not observe changes in genes involved in the innate immune response (**Figure 7**).

Genome editing strategies have also been developed by several groups including ours (Antoniani et al., 2018; Ramadier et al., 2021; Weber et al., 2020) with early, promising clinical results (Frangoul et al., 2020) but long-term efficacy and safety needs to be demonstrated, particularly in light of the recent findings on the CRISPR/Cas9-induced genotoxicity (Boutin et al., 2022; Nahmad et al., 2022). On the contrary, our approach can be readily implemented in clinically accepted LV design and eventually translated to the bedside after preclinical biosafety and biodistribution studies, to change the goal of current gene therapy clinical trial from an amelioration of clinical signs and reduction of blood transfusion to a complete cure of the disease.

## Acknowledgements

This work was supported by State funding from the Agence Nationale de la Recherche under “Investissements d’avenir” program (ANR-10-IAHU-01 and ANR-20-CE17-0016), the Paris Ile de France Region under “DIM Thérapie génique” initiative, and the Imagine Institute (Innogrant). We thank Christine Bole and the Imagine genomic facility for the generation of the NGS data.

## Author Contributions

MB designed, conducted experiments, analyzed data and wrote the paper, AC, PM, VP, SS, SR conducted experiments and analyzed data. GF, OR, FM, MC and MA contributed to the design of the experimental strategy. CM and OR analyzed NGS data. AM conceived the study, designed experiments and wrote the paper.

## Conflict of interest

MB, FM, MC, MA and AM are the inventors of two patents describing bifunctional LVs for hemoglobinopathies. All other authors declare no competing interests.

## Supplementary Figure Legends

**Figure S1:**

Infectious [Transduction Unit (TU)/ml] and physical titers of the bifunctional (βAS3/miRBCL11A-LVs) and control (βAS3-LV) vectors. Infectious titers were measured on HCT116 cells, 4 days after transduction (n=1-3 independent LV production).

**Figure S2:**

Gene transfer efficiency (VCN/cell) obtained for each MOI tested in HUDEP-2 cells transduced with the βAS3-, or βAS3m/miRBCL11A_Int2_del or _Int2 LV vectors. VCN/cell was measured 14 days after transduction.

**Figure S3:**

Mobilized peripheral blood HSPCs from two β-thalassemia donors (β^+^/β^+^ #1 and β^0^/β^0^ #2) were either not transduced (Mock; n=1 per donor) or transduced with control (βAS3 or βAS3/miRnt; n=4 independent biological replicates per donor) or bifunctional (βAS3/miRBCL11A; n=2 independent biological replicates per donor) vectors at different MOI for 24 hours. After transduction, cells were plated in clonogenic cultures to evaluate BFU-E and CFU-GM after 14 days, or differentiated in erythroid precursors. VCN/cell were measured 14 days after transduction by ddPCR in erythroblasts, BFU-E and CFU-GM pools or in BFU-E single colonies. βAS3- and βAS3/miRnt-transduced samples were pooled in all the analyses. **(A)** Average VCN/cell (mean±SD) in pools of BFU-E or CFU-GM, and in erythroblasts. **(B)** Frequency of BFU-E and CFU-GM in mock- and LV-transduced HSPCs. Results are represented as % of colonies obtained from 500 plated HSPCs and shown as mean±SD. **(C)** Clonal analysis of VCN/cell (mean±SD) in BFU-E single colonies (n=14 for βAS3 and 24 for βAS3/miRBCL11A).

**Figure S4:**

Expression of erythroid markers measured by flow cytometry along the differentiation (day 13, 16 and 20) in 2 β-thalassemia donors (n=1, 2, and 4 independent biological replicates per donor for Mock-, βAS3/miRBCL11A-, and βAS3- or βAS3m/miRnt-transduced samples, respectively). In the top panels, we plotted the frequencies of CD36^+^ (top left panel), CD71^+^ (top middle panel), and CD235a^+^ (top right panel) erythroid cells. During erythroid differentiation, cells progressively lose CD36 and CD71 expression. In the bottom panel, we plotted the frequencies of CD49^+^, Band3^+^ and CD49^+^Band3^+^ cells among the CD235a^+^ erythroid cells. During erythroid differentiation, CD235a^+^ cells lose progressively the CD49 marker and express Band3. βAS3- and βAS3m/miRnt-LVs: VCN/cell=1.2±0.3 and 2.0±0.2 for patient #1 and #2, respectively; βAS3/miRBCL11A-LV: VCN/cell=1.3±0.1 and 2.0±0.1 for patient #1 and #2, respectively.

**Figure S5:**

Expression of erythroid markers and enucleation rate measured by flow cytometry along the differentiation (day 13, 16 and 20) in 3 SCD donors (n=3, 7, and 10 independent biological replicates for mock-, βAS3- or βAS3/miRnt-, and βAS3/miRBCL11A-transduced samples, respectively). βAS3- and βAS3m/miRnt-LVs: VCN/cell=2.3±1.7; βAS3/miRBCL11A-LV: VCN/cell=1.8±0.7.

**Figure S6:**

Mobilized (2 donors) or non-mobilized (1 donor) peripheral blood HSPCs from three SCD donors were either not transduced (Mock; n=3 independent biological replicates) or transduced with control (βAS3 or βAS3m/miRnt; n=7 independent biological replicates) or bifunctional (βAS3/miRBCL11A; n=10 independent biological replicates) vectors at different MOI for 24 hours. After transduction, cells were plated in clonogenic cultures to evaluate BFU-E and CFU-GM after 14 days, or differentiated in erythroid precursors. VCN/cell were measured 14 days after transduction by ddPCR in erythroblasts and in BFU-E and CFU-GM pools. βAS3- and βAS3m/miRnt-transduced samples were pooled in all the analyses. **(A)** Frequency of BFU-E and CFU-GM in mock- and LV-transduced HSPCs. Results are represented as % of colonies obtained from 500 plated HSPCs and shown as mean±SD. ns, two-way ANOVA test. **(B)** Average VCN/cell (mean±SD) in pools of BFU-E or CFU-GM, and in erythroblasts. ns, two-way ANOVA test. **(C)** HbAS3 (left panel), HbF (middle panel) and HbS (right panel) expression measured by CE-HPLC in pools of BFU-E.

**Figure S7:**

Infectious [Transduction Unit (TU)/ml] and physical titers of βAS3m/miR7m and control (βAS3- and βAS3m/miRnt-LVs) vectors. Infectious titers were measured on HCT116 cells, 4 days after transduction. (n=3 independent LV production)

**Figure S8:**

(A,B) Mobilized (2 donors) or non-mobilized (1 donor) peripheral blood HSPCs from SCD donors were either mock-transduced (Mock; n=3 independent biological replicates) or transduced with control (βAS3 or βAS3m/miRnt; n=1-3 independent biological replicates per MOI) or bifunctional (βAS3m/miR7m; n=1-2 independent biological replicates per MOI) vectors at different MOI for 24 hours (n=2-3 per donor). After transduction, cells were plated in clonogenic cultures to evaluate BFU-E and CFU-GM after 14 days, or differentiated in erythroid precursors. VCN/cell were measured 14 days after transduction by ddPCR in erythroblasts, BFU-E and CFU-GM pools, and in BFU-E single colonies. βAS3- and βAS3m/miRnt-transduced samples were pooled in all the analyses. **(A)** BFU-E and CFU-GM frequencies evaluated in mock- and LV-transduced HSPCs. Results are represented as % of colonies obtained from 500 plated HSPCs, performed in duplicates and shown as mean±SD. ns, two-way ANOVA test. **(B)** Average VCN/cell (mean±SD) in bulk BFU-E, CFU-GM and erythroblasts. ns, two-way ANOVA test. **(C)** Analysis of VCN/cell in individual BFU-E (1 donor; n=1-2 independent biological replicates per MOI). Results are shown as mean±SD when applicable. The MOI is indicated below each graph. ns, two-way ANOVA test.

**Figure S9:**

**(A)** HbAS3 (left panel) and HbS (right panel) expression measured by CE-HPLC in pools of BFU-E (3 donors). Linear regression for HbAS3 and HbS expression, ***P<0.001. (n=3, 10, and 19 individual biological replicates for Mock-, βAS3m/miR7m, and βAS3- or βAS3m/miRnt-tranduced samples, respectively). **(B)** Analysis of β-like globin expression (RT-qPCR) in individual BFU-E derived from HD (1 donor) HSPCs transduced with βAS3- or βAS3m/miR7m-LV (n=14, and 24 individual BFU-E colony from samples transduced with βAS3- or βAS3m/miR7m-LV, respectively). Results are shown as mean±SD. The VCN/cell is indicated below each graph.

**Figure S10:**

Gene ontology analysis of LV- (βAS3 or βAS3m/miR7m) targeted genes defined by a read count > 95th percentile. BP, biological function (1 healthy donor; n=1 per LV).

## Notes

### Competing Interest Statement

The authors have declared no competing interest.

