## Supplemental Figure File for "Novel lentiviral vectors for gene therapy of sickle cell disease combining gene addition and gene silencing strategies"

### Figure S1, related to Figure 1

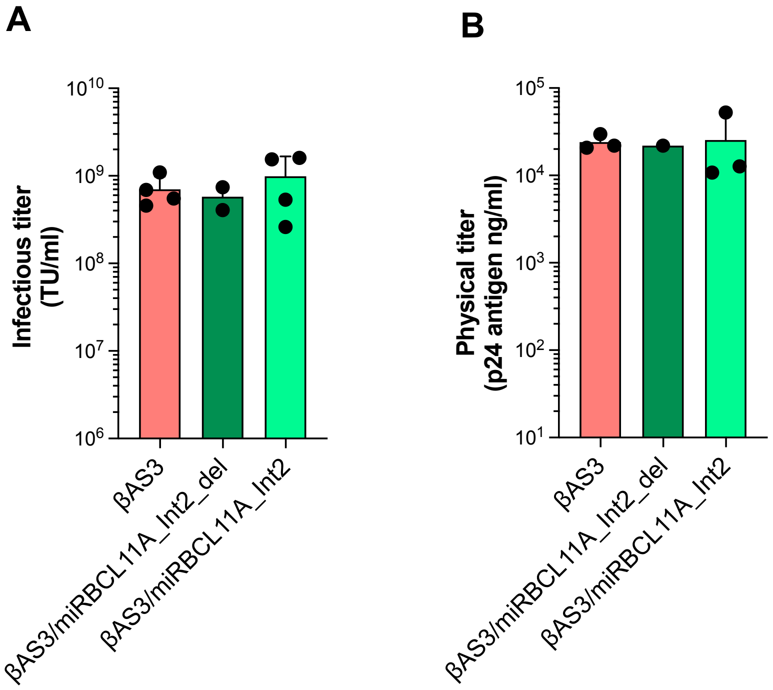

### Figure S2, related to Figure 1

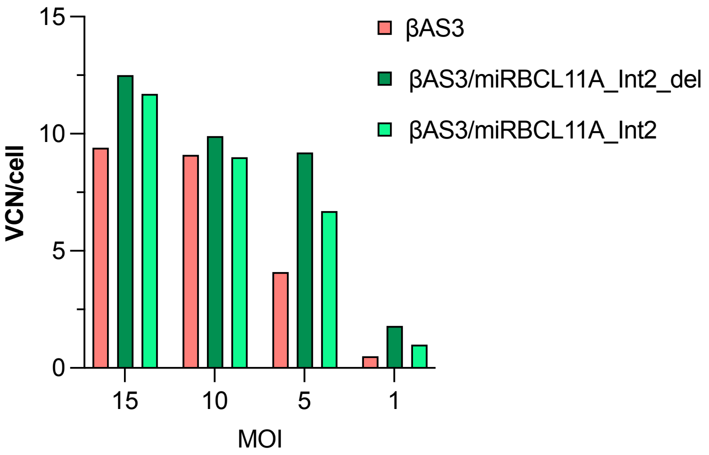

### Figure S3, related to Figure 2

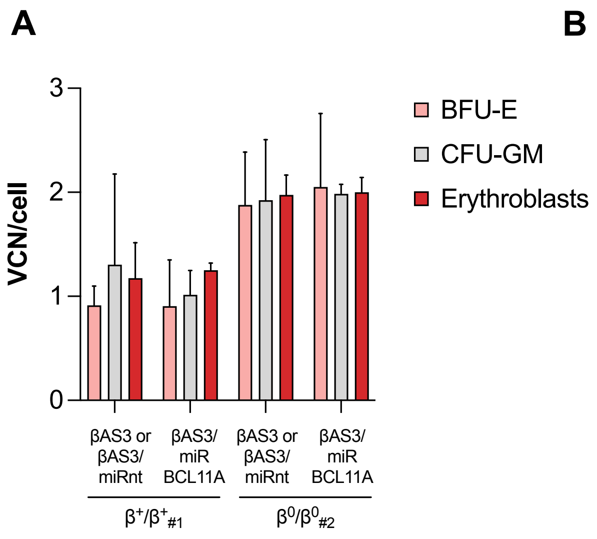


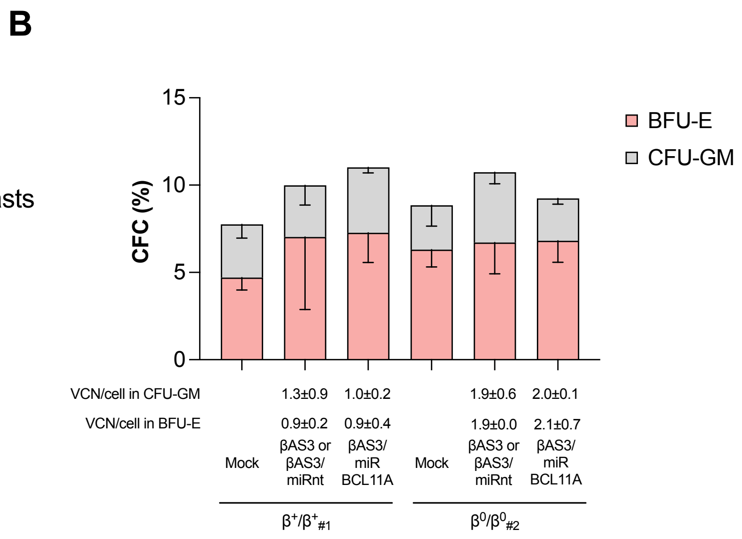


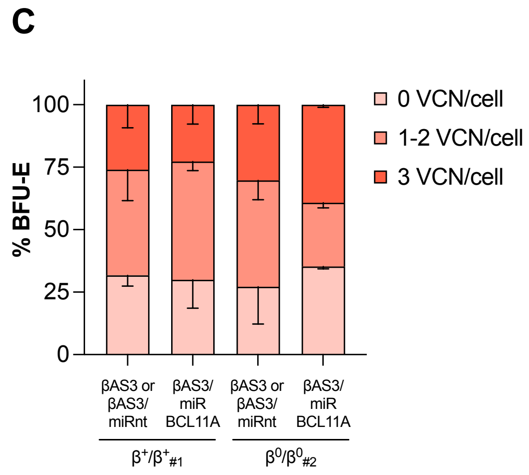

### Figure S4, related to Figure 2

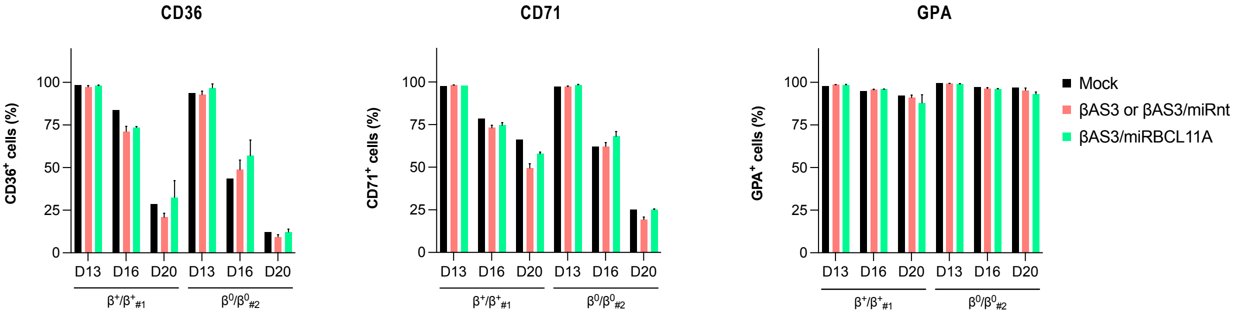


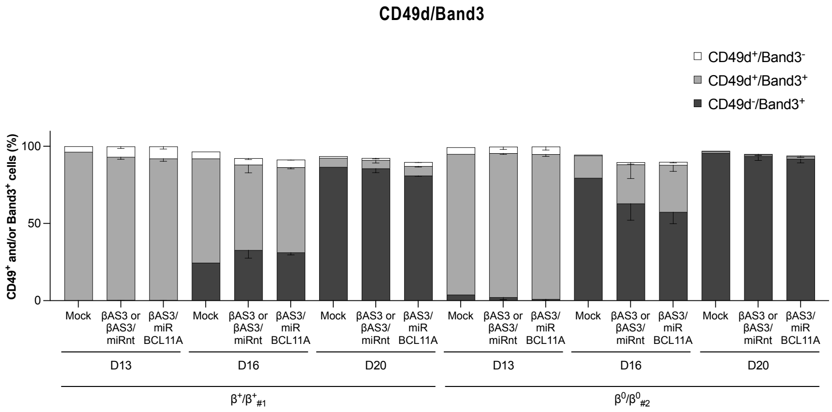

### Figure S5, related to Figure 3

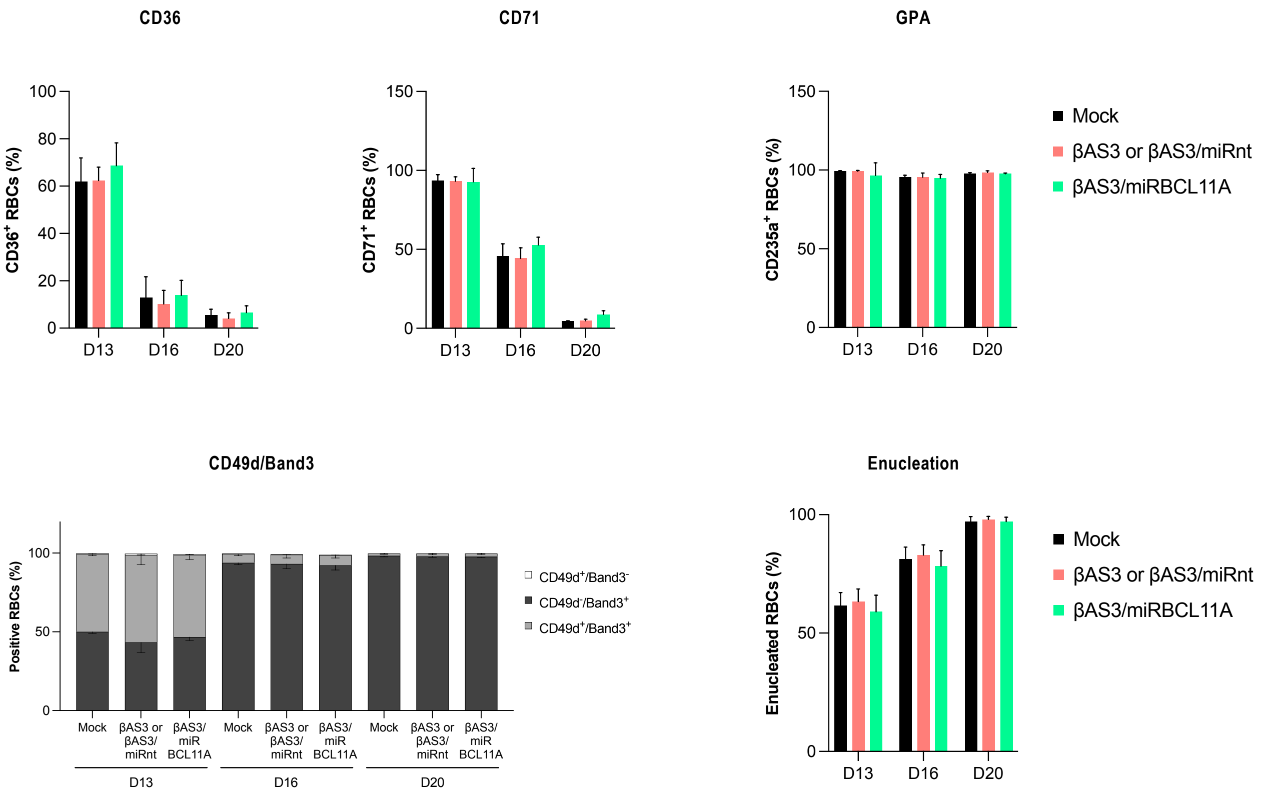

### Figure S6, related to Figure 3

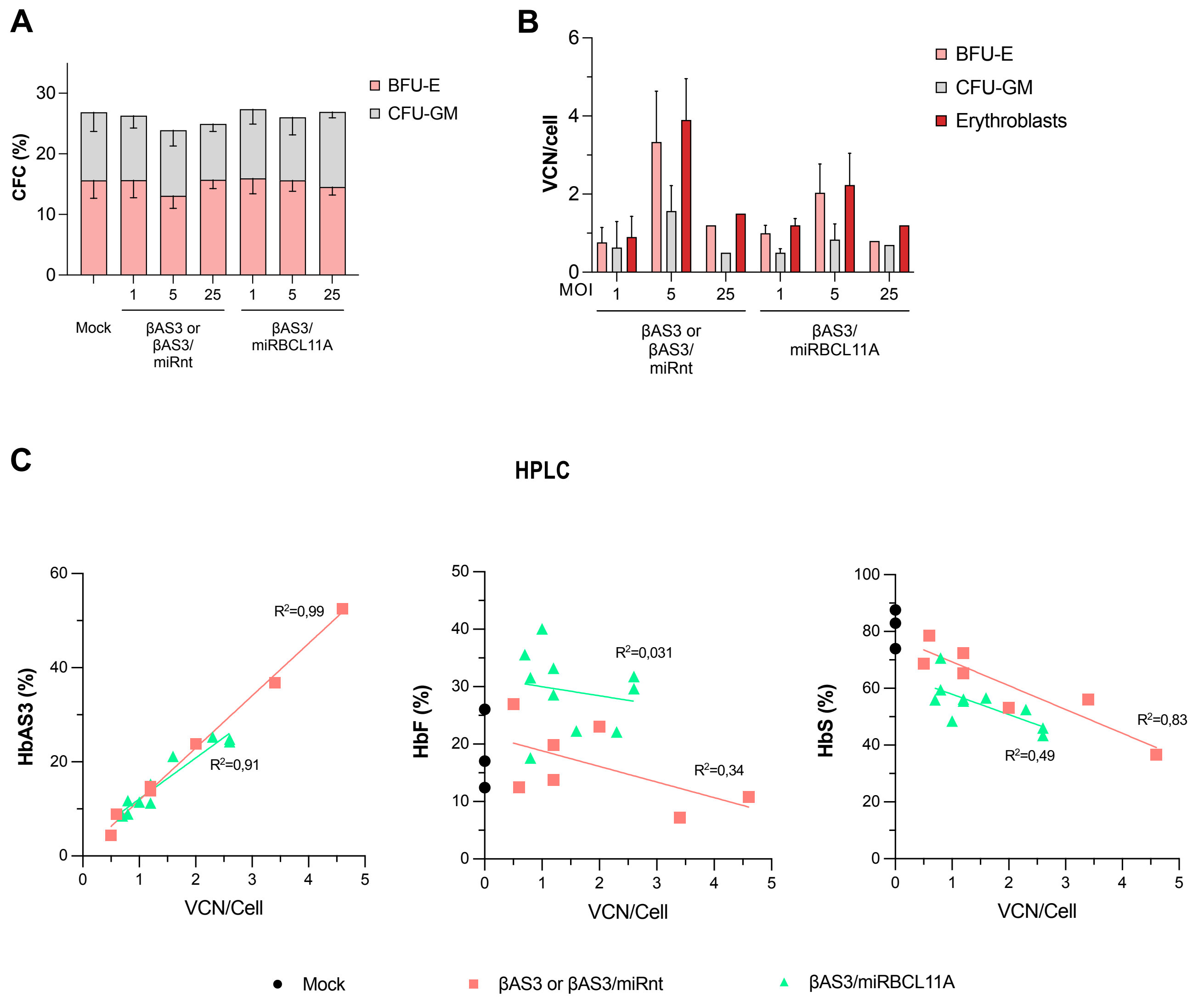

### Figure S7, related to Figure 5

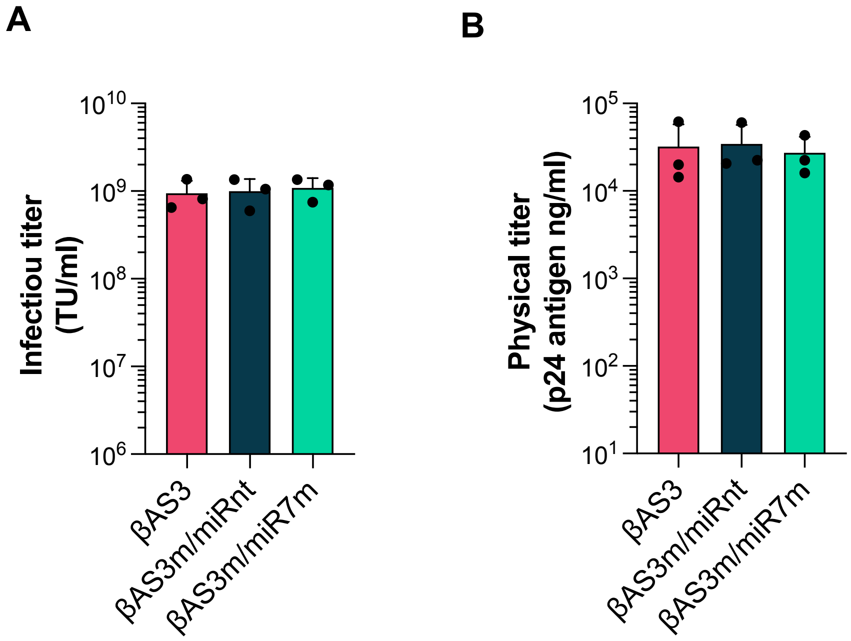

### Figure S8, related to Figure 5

A

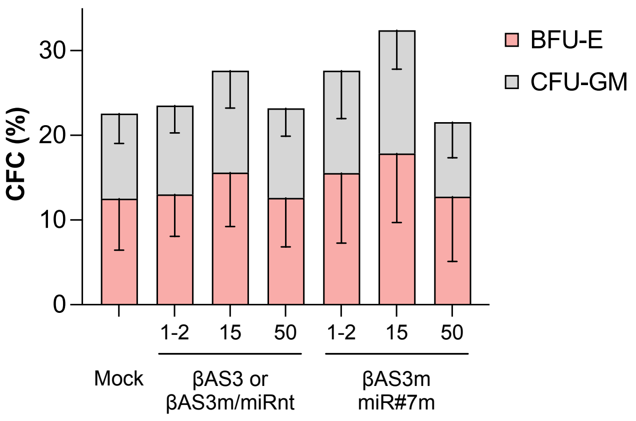


B

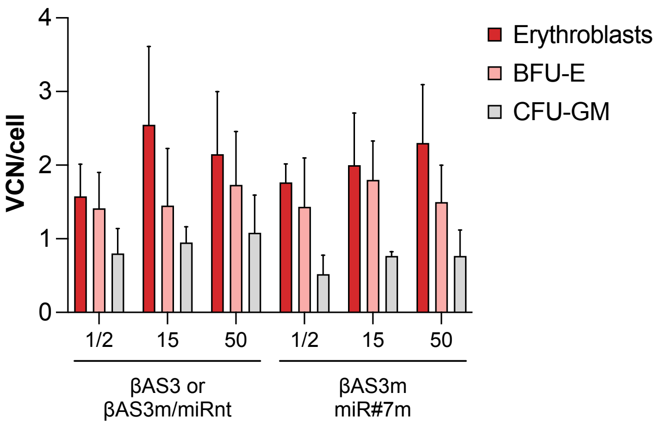


C

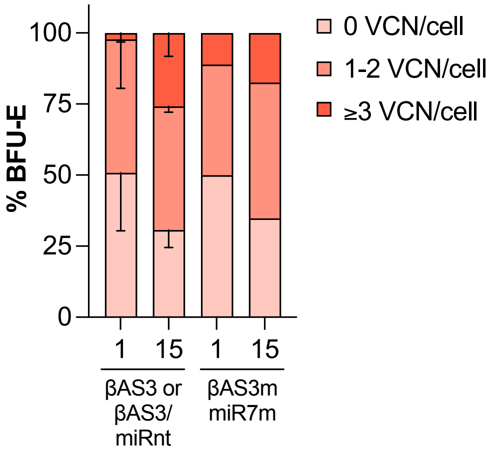

### Figure S9, related to Figure 5

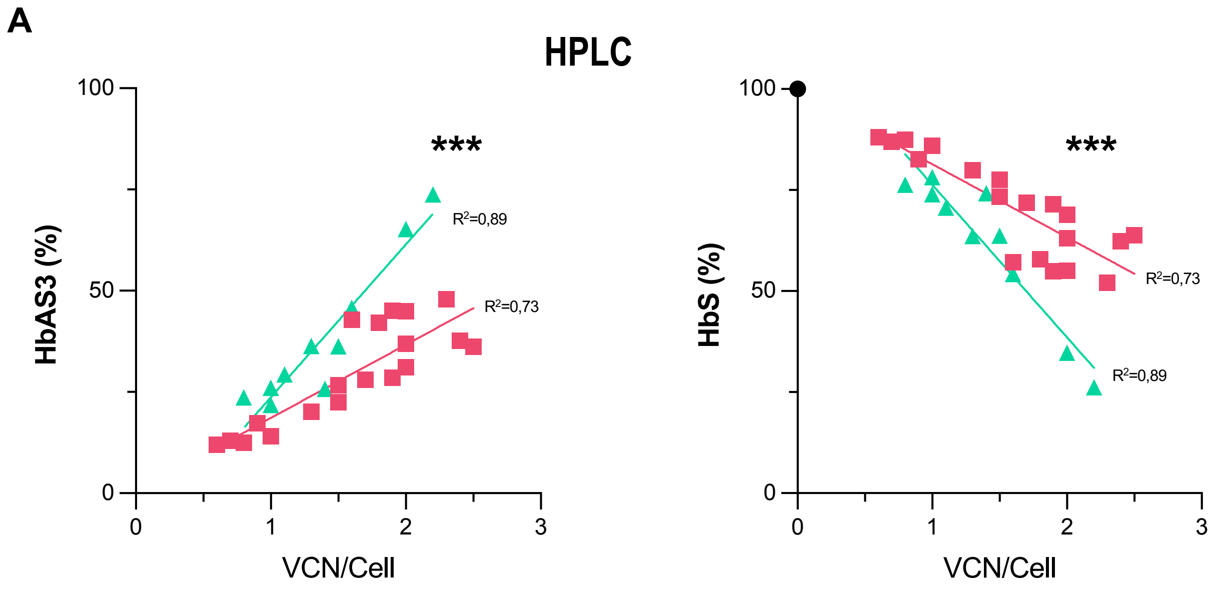


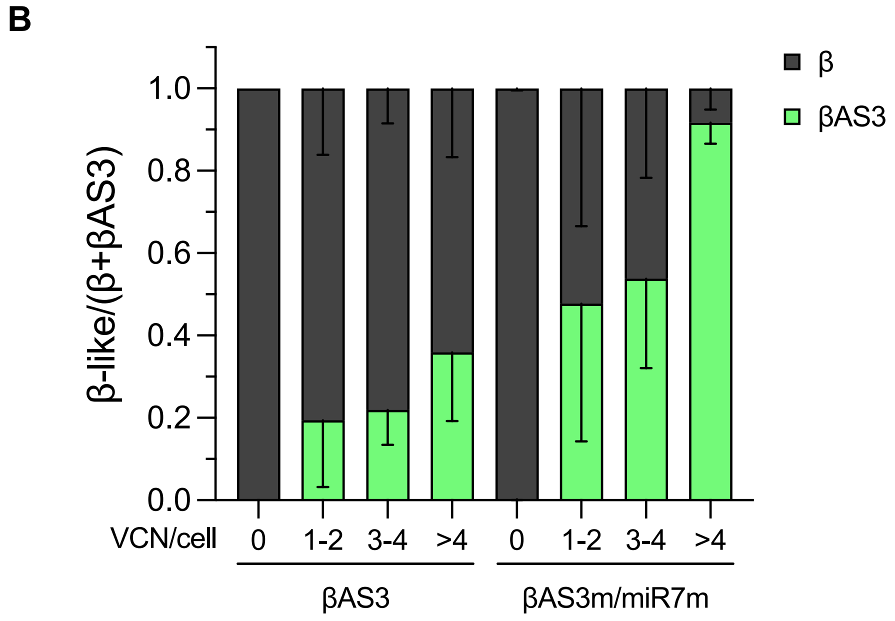

### Figure S10, related to Figure 7

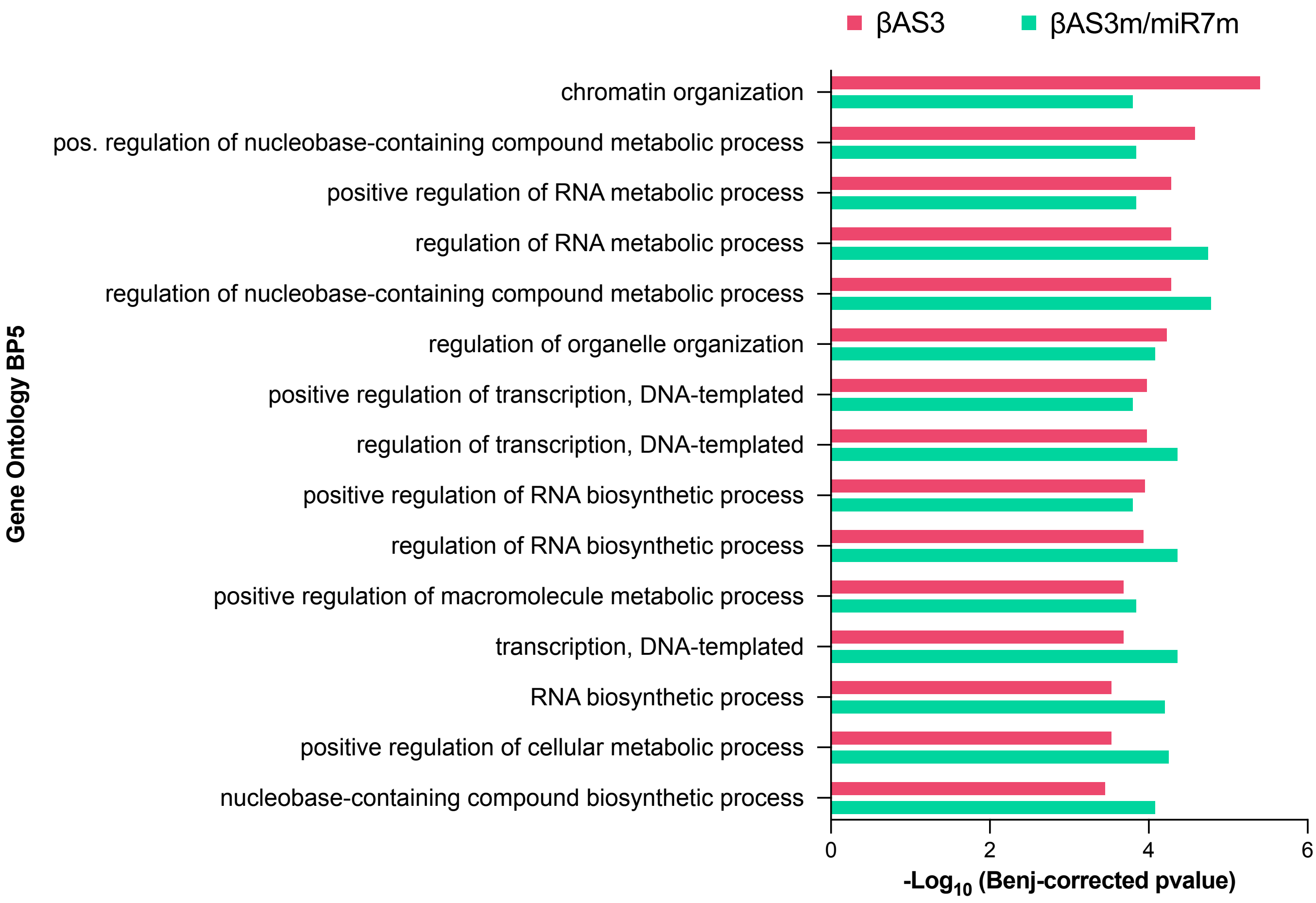
